# Intergroup lethal gang attacks do not require fission-fusion dynamics to evolve

**DOI:** 10.1101/2020.12.25.409938

**Authors:** Laura Martínez-Íñigo, Antje Engelhardt, Muhammad Agil, Malgorzata Pilot, Bonaventura Majolo

**Affiliations:** School of Psychology, University of Lincoln, UK; School of Life Sciences, University of Lincoln, UK; School of Biological and Environmental Science, Liverpool John Moores University, UK; Faculty of Veterinary Medicine, Bogor Agricultural University, Indonesia; Museum and Institute of Zoology, Polish Academy of Sciences

**Keywords:** between-group competition, coalitionary aggression, imbalance-of-power hypothesis, intergroup conflict, killing, raiding

## Abstract

Lethal gang attacks, in which multiple aggressors attack a single victim, are among the most widespread forms of violence between human groups. Chimpanzees (*Pan troglodytes*), as well as wolves (*Canis lupus*), spotted hyaenas (*Crocuta crocuta*), and lions (*Panthera leo*), perform gang attacks during raids. In raids, a few individuals of a group enter another group’s territory and attack its members if found in numerical disadvantage. Current theory predicts that raids and gang attacks are linked to fission-fusion dynamics, i.e., the capacity of a group to split into smaller subgroups of variable size and composition. However, over the last decade, research on social mammals without fission-fusion societies nor raiding have shown that they may also be involved in intergroup lethal gang attacks. Thus, neither fission-fusion dynamics nor raiding are required for gang attacks to evolve. Based on our first-ever reports of intergroup gang attacks in the crested macaque (*Macaca nigra*), combined with the synthesis of earlier observations of such attacks in several species living in stable groups, we develop a new hypothesis about the proximate causes leading to lethal intergroup aggression. We propose that the ability to estimate numerical odds, form coalitionary bonds, and show hostility towards outgroup individuals may suffice to trigger intergroup gang attacks when the conditions favour an imbalance of power between victims and attackers.

## INTRODUCTION

Violence between human groups has caused vast losses of lives through history and prehistory (Allen & Jones, 2014; Durrant, 2011; Glowacki et al., 2017; Kissel & Kim, 2019 but see Kelly, 2000). One of the most widespread forms of intergroup lethal and severe violence in humans are gang attacks during raids (Keeley, 1996; Otterbein, 2004). In raids, a group of men enters their enemies’ territory to find isolated or otherwise vulnerable individuals (Keeley, 1996). When they find them, multiple people concentrate physical aggression on a single victim (i.e. gang attack, Wilson et al., 2002), intending to kill it. Then, the attackers return to their territory (Keeley, 1996). Raids are the most widespread act of warfare among hunter-gatherers (Allen & Jones, 2014; Gat, 1999) and are a common tactic for guerrillas (Department of the Army, 1951) and youth gangs of industrialised societies (Wrangham & Wilson, 2004).

Raids similar to those of humans are frequent among common chimpanzees (*Pan troglodytes*). Sub-groups of chimpanzees, mostly males, enter their neighbours’ territory, apparently searching for members of other communities (Wrangham, 1999). If the raiding party detects individuals of a neighbouring community and has a numerical advantage over them, the raiding party attacks and often kills those individuals (Manson & Wrangham, 1991; Wrangham, 1999). In both humans and chimpanzees, gang attacks during raids seem prompted by a high numerical imbalance between aggressors and victims (Manson & Wrangham, 1991; Wrangham, 1999). Similarly, groups of wolves (*Canis lupus*), lions (*Panthera leo*) and spotted hyenas (*Crocuta crocuta*) attack and kill members of other groups during territorial activities (Manson & Wrangham, 1991; Wrangham, 1999). Besides their capacity for gang attacks and raiding, all these species have in common their capacity for forming coalitionary bonds and their groups with fission-fusion dynamics (Manson & Wrangham, 1991; Wrangham, 1999). Fission-fusion dynamics means that groups may sub-divide into smaller units of different sizes and composition for variable periods. Based on these observations, the imbalance-of-power hypothesis (Manson & Wrangham, 1991; Wrangham, 1999) postulates that intergroup lethal gang attacks occur in species with coalitionary bonds and fission-fusion dynamics. Coalitionary bonds enable gang attacks, while fission-fusion dynamics increase the chances of encounters with high numerical imbalance (Wrangham, 1999). Coalitionary bonds and fission-fusion dynamics together allow attacks at low cost for the aggressors, which may suffice to trigger lethal violence (Enquist & Leimar, 1990).

The imbalance-of-power hypothesis further predicts that attackers and victims would tend to be philopatric group members, i.e., individuals who remain in their natal group throughout their life (Wrangham, 1999). This prediction is related to the proposed ultimate function of intergroup killing; to increase intergroup dominance (“the intergroup dominance hypothesis”, Crofoot & Wrangham, 2010). Bigger groups tend to displace smaller groups (Majolo et al., 2020); thus, reducing the size of neighbouring groups would improve the odds of winning future encounters. Intergroup dominance improves access to fitness-enhancing resources and mating partners (Crofoot & Wrangham, 2010; Lemoine, Boesch, et al., 2020; Lemoine, Preis, et al., 2020; Mitani et al., 2010). On average, philopatric group members have more kin in their group than non-natal individuals (Chapais & Berman, 2004). Consequently, by participating in gang attacks, philopatric individuals have better chances to improve their inclusive fitness than non-natal members. Moreover, philopatric individuals are replaced by birth, which is slower than migration (Wrangham, 1999). Therefore, eliminating philopatric members of competing groups would produce a longer-lasting reduction in their group size than targeting non-natal individuals.

Contrary to the imbalance-of-power-hypothesis predictions, several mammal species with stable groups have been observed engaging in lethal gang attacks (Table 2 and Dyble et al., 2019; Johnstone et al., 2020). Here, we present a comprehensive and detailed account of intergroup gang attacks in a non-hominid primate species living in stable groups without fission-fusion dynamics, the crested macaque (*Macaca nigra*). We analyse 25 gang attacks in three groups of wild crested macaques, leading to at least six deaths over 13 years of observation. We test whether the sex of the attackers and victims of gang attacks was biased towards females, the philopatric sex. Such bias would be consistent with gang attacks having the ultimate function of increasing intergroup dominance. Based on our observations and those in other species, we propose an alternative hypothesis to explain the proximate causes of intergroup lethal gang attacks.

## METHODS

### Study Site and Subjects

Observations took place between 2006 and 2019 in the Tangkoko-Batuangus Nature Reserve (1°33’N, 125°10’E), in North Sulawesi, Indonesia. We studied three neighbouring groups of habituated crested macaques (R1, R2, and PB). Crested macaques, a critically endangered species endemic from the North Sulawesi (Supriatna & Andayani, 2008), live in groups with multiple adult females and males with their offspring. Groups typically contain more females than males and travel in cohesive units. Males usually abandon their natal group after adulthood and switch groups several times throughout their lifetime (Marty et al., 2016). Males may form coalitions (Neumann, 2013), but overall, they have weak bonds marked by avoidance (Tyrrell et al., 2020). Female crested macaques stay in their natal group and form coalitions between them, regardless of age, relatedness, or social bond strength (Duboscq, 2013). Home ranges of neighbouring crested macaque groups overlap almost entirely (Martínez-Íñigo, 2018; O’Brien & Kinnaird, 1997), and encounters between them take place several times per week ((Kerhoas et al., 2014; Martínez-Íñigo, 2018; O’Brien & Kinnaird, 1997)and Table S2 in ESM). Females participate aggressively during encounters, but male involvement is more common (Martínez-Íñigo, 2018).

R1 and R2 groups have been followed almost daily since 2006 and PB since 2008. At least three other groups are neighbours to R1, R2 and PB. One of these groups (R3), was partially habituated to the presence of researchers. The other neighbouring groups were unhabituated and fled when they detected people. In 2014, PB split into two groups (PB1 and PB2; only PB1 continued to be studied); in 2017 PB1 split into two groups (PB1A and PB1B; only PB1B continued to be studied). The study groups relied almost entirely on natural food apart for some sporadic raids to human resources (e.g. garbage). R2 and R1 often encountered small groups of tourists in the forest, whereas PB/PB1/PB1B had minimal contact with tourists. At the time of the recorded gang attacks, the average number of adults was 36.0 in R1 (average N of males=10.8; average N of females=25.2), 29.8 in R2 (males=6.9; females=22.9) and 26.9 in PB/PB1/PB1B (males=6.1; females=20.8). We could not identify sub-adults and juveniles individually, and we did not have reliable counts for these two age classes.

We defined females to be adults once they had given birth to an infant. We defined males as adults once their canines have fully erupted, and their testes had descended. Infants were monkeys who were ≤1 year old (age from birth records in habituated groups), or ventrally clinging to a female during encounters in non-habituated groups. Sub-adults displayed secondary sexual characters (e.g. sexual swelling in females, incipient canines in males) that were not fully developed as for the adult definitions above. All the other subjects fell into the juvenile class.

### Data Collection

We recorded intergroup gang attacks *ad libitum* (Altmann, 1974) employing videos, pictures and notes. We defined intergroup gang attacks as events in which two or more individuals of one group directed contact aggression against a conspecific from outside their group (Wilson et al., 2002). Contact aggression consisted of bites, hits and pulling from body parts. Individuals who hit, bit and/or pulled body parts of a victim during a gang attack were considered attackers. Individuals surrounding the victim without participating in the contact aggression were considered bystanders. If victims of gang attacks escaped and died within a week of the attacks, with no suspected cause of death other than the injuries inflicted during the attack, we considered that they had died due to the gang attack. We individually identified the individuals involved in gangs attacks whenever the visibility and the habituation allowed it.

### Data Analysis

We used a chi-square test to analyse whether the number of observed victims (or attackers) was sex-biased. Our study groups’ average sex ratios were 1:2.9 for victims’ groups; 1:2.5 for attackers’ groups. We used these sex ratios to calculate the expected values as follows. We divided the total number of adult males and females present in the group (T) by the sum of the sex ratio and multiplied this figure by the proportion of adults of each sex. When necessary, we converted the proportions to avoid non-integers (e.g., 1:2.9 turns into 10:29). For example, to calculate expected values for the attackers’ groups, we divided T by 7, which was the sum of 2 males per 5 females for a 1:2.5 sex ratio. Then we multiplied this figure by 2 to obtain the expected number of male attackers and by 5 for the expected number of female attackers. For some gang attacks, the number of adult attackers was reported as ≥2, without a specific figure (Table S1 in ESM). For those cases, we assumed the number of adults as 2 in the statistical analyses, to avoid inflating the number of attackers. We used R 3.4.2.(R Core Team 2020) to run the chi-square tests. No animal was the victim of more than one attack, but some adult females were involved in more than one attack. Unfortunately, for most gang attacks occurring before 2015, we had the total number of adult attackers but not their ID. Thus, the number of adult attackers entered in the chi-square test is at risk of being inflated, if a few adult attackers took part in more than one case of intergroup violence. To control for this possibility, for each case of intergroup violence we divided the number of adult attackers of each sex by the total number of adult males/females living in the attackers’ group at the time of the recorded case of intergroup violence. We entered these proportions of adult attackers of each sex as the response variable in a generalised linear mixed model (GLMM) run using Stata v.12.1(StataCorp 2011). Sex of the attacker was the fixed factor, and we entered group ID as a random factor.

### Ethical note

Our research protocol adhered to standards as defined by the Ethics Committee of the University of Lincoln and the principles of “Ethical Treatment of Non-Human Primates” as stated by the American Society of Primatologists. Our research adhered to the legal requirements of the Indonesian and UK governments. Permission to conduct the study in the field (229/SIP/FRP/SM/VIII/2015) was granted by the Indonesian State Ministry of Research and Technology (RISTEK).

## RESULTS

### Gang Attacks

Between March 2006 and May 2019, we documented 25 intergroup gang attacks (Table S1 in ESM). Based on the seven best-documented cases (see those of 2015-2016 in ESM), gang attacks often happened during intergroup encounters in which the attacking group surrounded an individual of the opposing group during a chase. The group companions of the victim retreated, leaving it isolated. From those individuals of the opposing group surrounding the target, a few attacked the victim (hereafter, attackers) while others observed the attack without intervening (hereafter, bystanders). Most members of the attacking group continued with their regular activities and were considered neither attackers nor bystanders.

Most gang attacks (N=21; 84%, Table 1) occurred during intergroup encounters. Two additional attacks (8%) targeted adult males travelling alone. The context of the remaining two attacks (8%) is unknown but suspected to have occurred during intergroup encounters.

**Table 1.** The number of victims and attackers involved in gang attacks per group and age-sex class.

| Victims<sup>1</sup> |  | Attackers<sup>2</sup> |  | Gang attack |  |
| --- | --- | --- | --- | --- | --- |
| Age-sex class | Group | Age-sex class | Group | Scenario<sup>3</sup> | Outcome<sup>4</sup> |
| Adult females: 9 | R1: 3 | Adult females: $\geq 17$ | R1: 20 | Intergroup encounters: 21 | Escaped: 19 |
| Adult females + infant: 6 | R2: 8 (+1?) | Adult males: $\geq 2$ | R2: 3 | Male travelling alone: 2 | Died in the attack: 1 |
| Adult males: 2 | PB/PB1/PB1A: 7 | Sub-adult females: $\geq 7$ | PB/PB1/PB1A: 2 | Unknown: 2 | Died after the attack due to injuries: 4 |
| Juveniles: 7 | R3: 1 (+1?) | Sub-adult males: $\geq 15$ | | | Died after the attack due to starvation/thirst: 1 |
| Sub-adult female: 1 | Unhabituated: 4 | Juveniles: $\geq 16$ | | | Unknown: 6 |

### Victims

Each gang attack targeted a single victim (N=19), except when the targets were females with unweaned infants (N=6). In total, 31 animals were the target of gang attacks. Adult females were the most frequent victim (Table 1), but there was no significant sex bias for victims when the sex ratio of their group was considered (*χ^2^_1_*=1.72, P=0.19). Six victims died because of the attacks (16.4% of victims; Table 1). Four of these victims were infants, and two were adult females. Three infants died from the injuries received during the attack; one infant was not injured but died from starvation after losing contact from its mother, who escaped the attack (see ESM).

### Attackers

The mean ratio of individually recognisable attackers per victim was 3.76 ±1.54 (X+SE, N= 6; these figures exclude infant victims). No wounds were observed on the attackers. 94.7% of adult attackers were females (Table 1). The bias in the attacker’s sex towards females was significant when the sex ratio was considered in the chi-squared test (χ^2^_1_=7.05, P<0.01) and the GLMM (Wald χ^2^= 19.14, coefficient ± SE = −0.06 ± 0.01, z = −4.37, P<0.001).

## DISCUSSION

Our long-term data showed that intergroup gang attacks in crested macaques fit several postulates of the imbalance-of-power hypothesis and differ in some key features (Manson & Wrangham, 1991; Wrangham, 1999). As predicted by the imbalance-of-power hypothesis, in intergroup gang attacks in crested macaques: 1) attackers outnumber victims; 2) the risk of injuries for the attackers is low, and 3) the philopatric sex is the most frequent attacker and victim. Contrary to the imbalance-of-power hypothesis, 1) crested macaques live in groups without fission-fusion dynamics, and 2) gang attacks mostly occur during intergroup encounters rather than during raids.

Our findings support the central prediction of the imbalance-of-power hypothesis; intergroup gang attacks are more likely when attackers’ costs are meager due to highly uneven numbers between attackers and victims (Manson & Wrangham, 1991; Wrangham, 1999). In addition to crested macaques, other primates (Table 2) and mammals (e.g. banded mongooses, *Mungos mungos*, Johnstone et al., 2020; and meerkats, *Suricata suricatta*, Dyble et al., 2019) living in cohesive groups, participate in intergroup lethal gang attacks. In all cases, attackers outnumber the victims and are rarely injured. Based on these observations, we argue that gang attacks have evolved independently from fission-fusion dynamics, contradicting the current version of the imbalance-of-power hypothesis. Intergroup gang attacks might result from the widespread animal ability to assess numerical odds (Benson-Amram et al., 2018; Furrer et al., 2011; Radford, 2003; Scarry, 2020; Tanner, 2006), coupled with coalitionary bonds and intergroup hostility. These features would suffice to trigger lethal and severe gang attacks whenever specific intergroup encounter conditions enabled attacks at low cost.

**Table 2.**
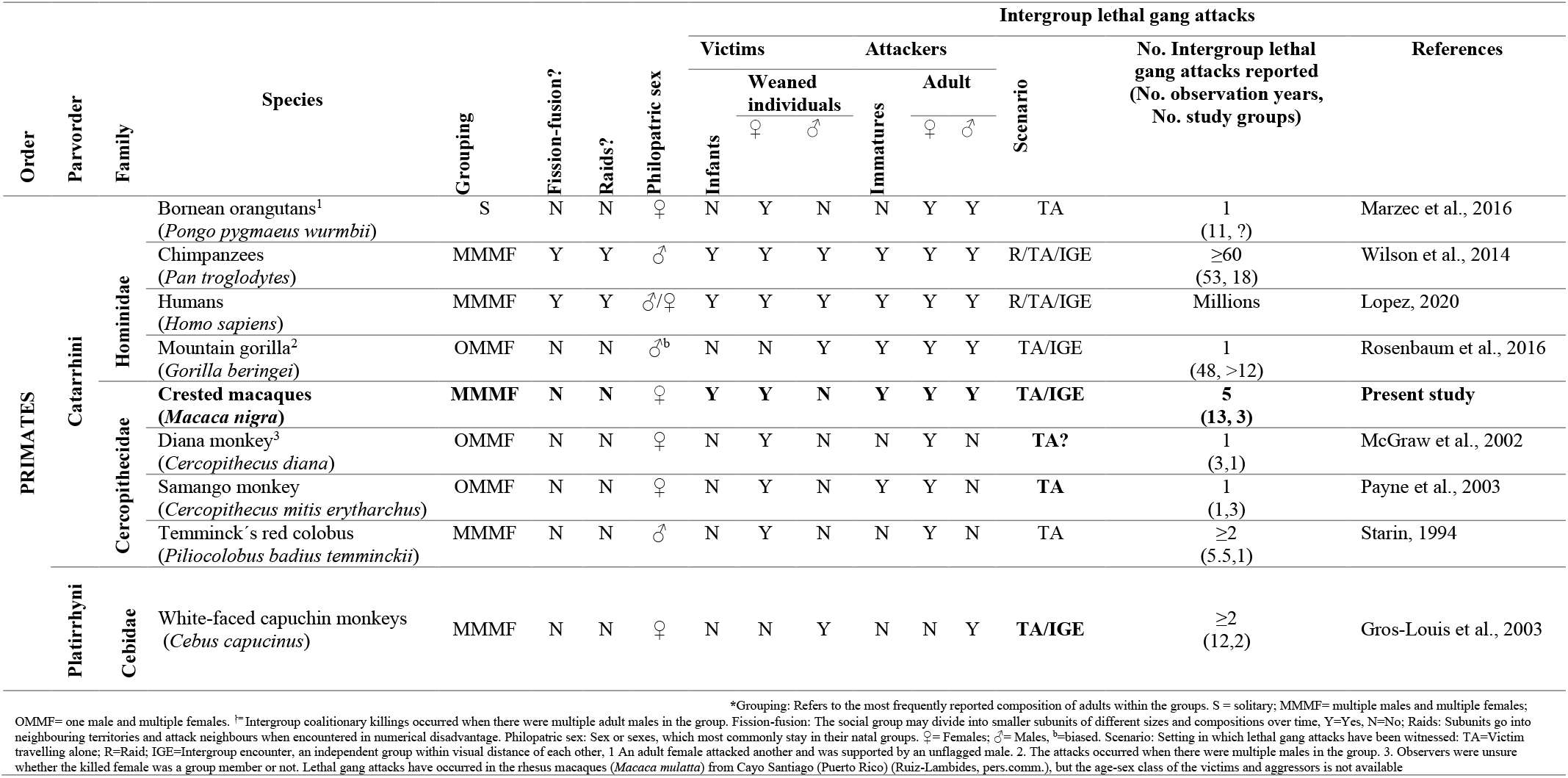
Summary of documented cases of intergroup lethal gang attacks in primates.

The conditions leading to intergroup gang attacks differ between species living in fission-fusion societies and species living in cohesive groups. In species with fission-fusion dynamics, such as spider monkeys (*Ateles geoffroyi yucatanensis*) (Aureli et al., 2006), intergroup gang attacks are common during raids, which significantly increase the opportunities for encounters between uneven numbers of opponents. Raids have not been observed in animals forming cohesive groups, as far as we are aware. Instead, intergroup gang attacks in species living in stable groups usually occur in two scenarios (Table 2): 1) A group finds an outgroup conspecific travelling alone; 2) A group isolates a member of another group during an intergroup encounter. The frequency of these two scenarios is likely to differ between species and populations, which could explain why many species with coalitionary bonds, intergroup hostility and the ability to assess numerical odds, have not been observed engaging in gang attacks against outgroup members.

On the one hand, individuals of cohesive groups may travel alone when searching for other groups to immigrate (Sue Boinski et al., 2005; Doolan & Macdonald, 1996; Janmaat et al., 2009; Sprague et al., 1998), which would increase their chances of suffering a gang attack. However, most species have strategies to avoid this lone phase, such as forming bachelor groups (Sprague et al., 1998; Sussman, 1992; Yao et al., 2011), immigrating in parallel with peers or kin (Schoof et al., 2009), and dispersing during intergroup encounters (Cheney & Seyfarth, 1983; Melnick et al., 1984; Yao et al., 2011). The more frequent these strategies are in a population, the less likely it will be for stable groups to find lone individuals.

On the other hand, chances of isolating individuals during encounters may depend on multiple factors such as the frequency of encounters themselves. For instance, crested macaques encounter their neighbours several times per week (Kerhoas et al., 2014; Kinnaird & O’Brien, 2000; Martínez-Íñigo, 2018) (see Table S2 in ESM), while most other primates do so once or less per month (Crofoot & Wrangham, 2010). Preliminary evidence in crested macaques suggests that a higher rate of intergroup encounters is associated with a greater rate of intergroup gang attacks when comparing between groups and within the same group over time (see ESM). Such results suggest that intergroup gang attacks’ rate may, indeed, be positively correlated to intergroup encounter rate.

Another factor that may increase the chances of isolating individuals during encounters is the habitat. In our study, gang attacks appeared to be more common when intergroup encounters happened in areas where natural barriers, such as the sea or dense vegetation, reduced the number of escape routes. Open habitats may facilitate potential victims to flee more readily than dense vegetation or environments with obstacles.

The primary function of intergroup gang attacks could differ between species living in stable groups and groups displaying fission-fusion dynamics. In cohesive species, attackers often target outsiders who otherwise could immigrate into the attacking group, threatening the attackers’ social rank and offspring (Gros-Louis et al., 2003; Rosenbaum et al., 2016; Starin, 1994). Other times, attackers aim for outgroup individuals to prevent intergroup mating during encounters (Johnstone et al., 2020). Therefore, victims of gang attacks in cohesive species often pose a direct threat to the attackers’ fitness. By engaging in the aggressions, attackers defend their offspring, immediate access to within-group mating partners, and/or their social status. This is not the case in fission-fusion species, where the principal targets and participants of gang attacks belong to the philopatric sex (Wrangham, 1999). Permanent residents of neighbouring groups affect each other’s fitness chiefly through between-group competition for resources and territory, which can, in turn, influence mate attraction and retention. Reducing the size of neighbouring groups is expected to improve access to such fitness-enhancing resources since larger groups tend to displace smaller ones (Crofoot & Wrangham, 2010; Lemoine, Boesch, et al., 2020; Lemoine, Preis, et al., 2020; Mitani et al., 2010). Killing philopatric members of other groups is the most cost-effective way of reducing the size of neighbouring groups, given that their replacement is slower than that of immigrants (Wrangham, 1999). In addition, group members other than the attackers profit from the benefits derived from gang attacks. The more relatives an attacker has within its group, the greater its inclusive fitness benefits would be. Since, on average, philopatric individuals, have more relatives in their group than immigrants have (Chapais & Berman, 2004), they have more to gain from intergroup gang attacks. In summary, participants of gang attacks in fission-fusion species benefit from the attacks through a potential increase in intergroup dominance, which improves their inclusive fitness.

Nevertheless, species living in stable groups may also use intergroup gang attacks as a strategy to increase their dominance over other groups. Aggressors and victims in crested macaques are commonly females, the philopatric sex. The prevalence of adult females as victims was non-significant when we controlled for sex ratio, but the two killed adults were females. Furthermore, the intergroup killing of infants in contexts that do not increase aggressors’ mating opportunities might be a strategy to defend long-term access to resources (Agoramoorthy & Rudran, 1995; Brown, 2020). In crested macaques, infants were the most common victim after adult females, and the most frequently killed. Intergroup infant killing during gang attacks is also common in meerkats (Dyble et al., 2019). In both, meerkats and crested macaques, larger groups tend to win intergroup encounters, with the number of pups and females, respectively, increasing the odds of displacing other groups (Dyble et al., 2019; Martínez-Íñigo, 2018). Samango monkeys (*Cercopithecus mitis erytharchus*) (Payne et al., 2003), and possibly Diana monkeys (*Cercopithecus diana*) (McGraw et al., 2002) are two other species in which philopatric individuals have gang attacked an outgroup individual of their sex. These patterns suggest that some species living in stable groups may use intergroup gang attacks to increase their dominance over other groups. Further research will be needed to confirm that such attacks have the expected impact on between-group competition in these species.

## Supporting information

ESM

## ACKNOWLEDGEMENTS

We gratefully acknowledge the permission of the Indonesian State Ministry of Research and Technology, the Directorate General of Forest Protection and Nature Conservation in Jakarta and the Department for the Conservation of Natural Resources in Manado, to conduct this research in the Tangkoko-Batuangus Nature Reserve. We are grateful to all the researchers and field assistants of the Macaca Nigra Project. We are particularly in debt with Rismayanti, Eka Cahyaningrum, Juliette M. Berthier, Meldy Tamengge, Dwi Yandhi Febriyanti, Iwan Halir and Julianus Mendome for their crucial contribution to the long-term data collection on intergroup violence. We thank Daphne Kerhoas and Margaret F. Kinnaird for their comments and data. We thank three anonymous reviewers who helped improve an earlier version of this manuscript and the crowdfunding contributors that helped fund this project. This work was supported by a RIF scholarship scheme (RIF2014S-32), from the University of Lincoln, awarded to LMI. MP was supported by the Polish National Agency for Academic Exchange (NAWA; Polish Returns Fellowship PPN/PPO/2018/1/00037).

