## Supplementary material for "Intergroup lethal gang attacks do not require fission-fusion dynamics to evolve": ESM

### Electronic Supplementary Material

**Pre-print server:** bioRxiv

#### Summary table on gang attacks in crested macaques

**Table S1**. Summary of intergroup gang attacks in three groups of crested macaques from Tangkoko Nature Reserve (North Sulawesi, Indonesia) recorded between 2006-2019

| Date | Victim | | Attackers | | Attack | | |
|---|---|---|---|---|---|---|---|
| | **Age-sex class** | **Group** | **Age-sex class (number)** | **Group** | **Scenario** | **Duration (min)** | **Outcome** |
| 03/08/2006 | Adult female | R2 | Adult females (x), Sub-adult males (x), Juveniles (x) | R1 | Intergroup encounter | ≈ 45 | Escaped wounded[α] |
| 15/11/2006 | Adult female | R2 | Unknown | R1 | Intergroup encounter | Unknown | Disappeared after encounter Body found days later |
| 2007 | Adult female | R1 | Adult females (x), Sub-adult males (x) | R2 | Intergroup encounter | Unknown | Escaped |
| 18/07/2007 | Adult female + Infant female | R2 | Unknown | R1 | Intergroup encounter | >2 | Female escaped wounded[α] with her infant |
| 08/10/2007 | Adult male | R1 | Unknown | R2 | Travelling alone | >49 | Escaped wounded[α] and disappeared |

NH=Non-habituated group; x ≥ 2 individuals (exact number unknown). Dates are given in the format dd/mm/yyyy, except when month and/or day were unknown. [α]Wounded/unwounded categories were assigned by observers from at least 5m away from the victims. This means that only major wounds could be detected.

**Table S1. (Cont. I)**

| Date | Victim | | Attackers | | Attack | | |
|---|---|---|---|---|---|---|---|
| | **Age-sex class** | **Group** | **Age-sex class (number)** | **Group** | **Scenario** | **Duration (min)** | **Outcome** |
| 14/11/2007 | Adult female | NH | Adult females (x), Sub-adult males (x), Juveniles (x) | R2 | Intergroup encounter | Unknown | Escaped wounded |
| 14/11/2007 | Adult female + Infant female | R2 | Adult females (1), Adult males (1), Sub-adult males (1), Juveniles (x) | R1 | Intergroup encounter | Unknown | Adult female escaped. Infant found dead one day later. The infant had wounds from an encounter before the gang attack |
| 16/02/2008 | Adult female + Infant | NH | Adult females (x), Sub-adult males (x), Juveniles (x) | R1 | Intergroup encounter | Unknown | Female escaped wounded. Infant was left in R1, where it was carried by a juvenile female. No further information available. |
| 16/06/2008 | Adult female + Infant male | R2 | Adult female: Adult males (1)<br><br>Infant: Adult females (1), Juveniles (x) | R1 | Intergroup encounter | Unknown | Female escaped unwounded with her wounded infant. He died 5 days later. |
| 02/11/2008 | Juvenile | R2 or R3 | Unknown | R1 | Intergroup encounter | >6 | Escaped wounded |

NH=Non-habituated group; x ≥ 2 individuals (exact number unknown). Dates are given in the format dd/mm/yyyy, except when month and/or day were unknown. [α]Wounded/unwounded categories were assigned by observers from at least 5m away from the victims. This means that only major wounds could be detected.

**Table S1. (Cont. II)**

| Date | Victim | | Attackers | | Attack | | |
|---|---|---|---|---|---|---|---|
| | **Age-sex class** | **Group** | **Age-sex class (number)** | **Group** | **Scenario** | **Duration (min)** | **Outcome** |
| 01/12/2008 | Juvenile | R2 | Adult females (4), Juveniles (x) | R1 | Intergroup encounter | ≈ 15 | Escaped unwounded |
| 20/12/2008 | Juvenile | PB | Adult females (3), Juveniles (x) | R1 | Intergroup encounter | Unknown | Unknown |
| 05/02/2009 | Adult female | PB | Adult females (3), Sub-adult males (1), Juveniles (x) | R1 | Intergroup encounter | Unknown | Escaped |
| 31/05/2009 | Adult female | R3 | Adult females (4), Sub-adult males (x) | R1 | Intergroup encounter | Unknown | Unknown |
| 24/11/2009 | Juvenile | R1 | Juveniles (x) | PB | Intergroup encounter | Unknown | Unknown |
| 01/01/2010 | Adult male | R2 | Unknown | R1 | Traveling alone | Unknown | Escaped wounded |
| 09/11/2010 | Juvenile | R3 | Unknown | R1 | Unknown | >120 | Unknown |
| 18/12/2015 | Adult female | NH | Adult females (1), Sub-adult females (1), Juveniles (x) | PB1 | Unknown | >27 | Died during attack |

NH=Non-habituated group; x ≥ 2 individuals (exact number unknown). Dates are given in the format dd/mm/yyyy, except when month and/or day were unknown. [α]Wounded/unwounded categories were assigned by observers from at least 5m away from the victims. This means that only major wounds could be detected.

**Table S1. (Cont. III)**

| Date | Victim | | Attackers | | Attack | | |
|---|---|---|---|---|---|---|---|
| | **Age-sex class** | **Group** | **Age-sex class (number)** | **Group** | **Scenario** | **Duration (min)** | **Outcome** |
| 20/02/2016 | Juvenile female | R2 | Sub-adult females (2), Sub-adult males (1), Juveniles (x) | R1 | Intergroup encounter | >48 | Escaped |
| 29/04/2016 | Adult female | NH | Adult females (2), Sub-adult males (1), Sub-adult females (3), Juveniles (x) | R1 | Intergroup encounter | >47 | Escaped |
| 18/07/2016 | Sub-adult female | PB1 | Adult females (1), Sub-adult females (1), Sub-adult males (2), Juveniles (x) | R1 | Intergroup encounter | >10 | Escaped unwounded |
| 18/07/2016 | Adult female | PB1 | Adult females (2), Sub-adult females (1), Sub-adult males (1) | R1 | Intergroup encounter | >3 | Escaped unwounded |

NH=Non-habituated group; x ≥ 2 individuals (exact number unknown). Dates are given in the format dd/mm/yyyy, except when month and/or day were unknown. [α]Wounded/unwounded categories were assigned by observers from at least 5m away from the victims. This means that only major wounds could be detected.

**Table S1. (Cont. IV)**

| Date | Victim | | Attackers | | Attack | | |
|---|---|---|---|---|---|---|---|
| | Age-sex class | Group | Age-sex class (number) | Group | Scenario | Duration (min) | Outcome |
| 18/07/2016 | Adult female + Infant female | PB1 | Adult female:<br>Adult females (3), Sub-adult females (2), Sub-adult males (2), Juveniles (x).<br><br>Infant female:<br>Adult females (1), Sub-adult males (1), Juveniles (x) | R1 | Intergroup encounter | Adult female:<br>>11<br><br>Infant:<br>>18 | Female escaped unwounded. Infant stayed in R1 and was carried by an adult female. The infant died the next day, presumably from thirst. |
| 18/07/2016 | Juvenile female | PB1 | Adult females (2), Sub-adult females (1), Sub-adult males (1), Juveniles (x) | R1 | Intergroup encounter | >4 | Escaped unwounded |
| 19/01/2019 | Adult female + Infant male | PB1B | Sub-adult males (4) | R1 | Intergroup encounter | Unknown | Female escaped. Wounded infant died 2h after the attack. |

NH=Non-habituated group; x ≥ 2 individuals (exact number unknown). Dates are given in the format dd/mm/yyyy, except when month and/or day were unknown. [α]Wounded/unwounded categories were assigned by observers from at least 5m away from the victims. This means that only major wounds could be detected.

#### Details on gang attacks in crested macaques

In the following pages, we provide the available evidence for each intergroup gang attack summarized in Table S1. Gang attacks were named as **harassment** and coded as "ar" in the long-term database of the Macaca Nigra Project (MNP). MNP defined harassment as: "*event in which several individuals threat, chase, bite, hit and grab together another one individual. Often during intergroup encounters, one individual of one group is harassed by many of the other.*"

#### 3rd August 2006: R1 harassed an adult female from R2 (Amber, AD)

**Available information about the attack:** 5 pictures taken by Antje Engelhardt during the harassment and field notes

**Figure S1.1:** 8:07- Amber, an adult female from R2, is curled up on the ground with two individuals sitting next to her. The individuals likely are a juvenile and an adult female. A third individual stands by Amber, but only some fingers are visible.

**Figure S1.2.:** 8:48-The face of Amber during the harassment. She is resting the left side of her face on the ground, and her eyes are half-open, giving the impression of being dead. She has an injury of a couple of centimetres on the right side of her head. The skin has been removed but is not bleeding.

**Figure S1.3**.: 8:50- Amber is facing down on the ground with three individuals around her, but only their arms are visible. None seems to have adult male size.

**Figure S1.4**.: 8:50 - Amber is in the same posture as in ***Error! Reference source not found.***.3., and with the same number of individuals, but one of them is inspecting her anal area.

**Figure S1.5**: 8:51- A sub-adult male sits by Amber while she lies down on the ground looking dead. There are at least two other individuals around her.

*Pictures*

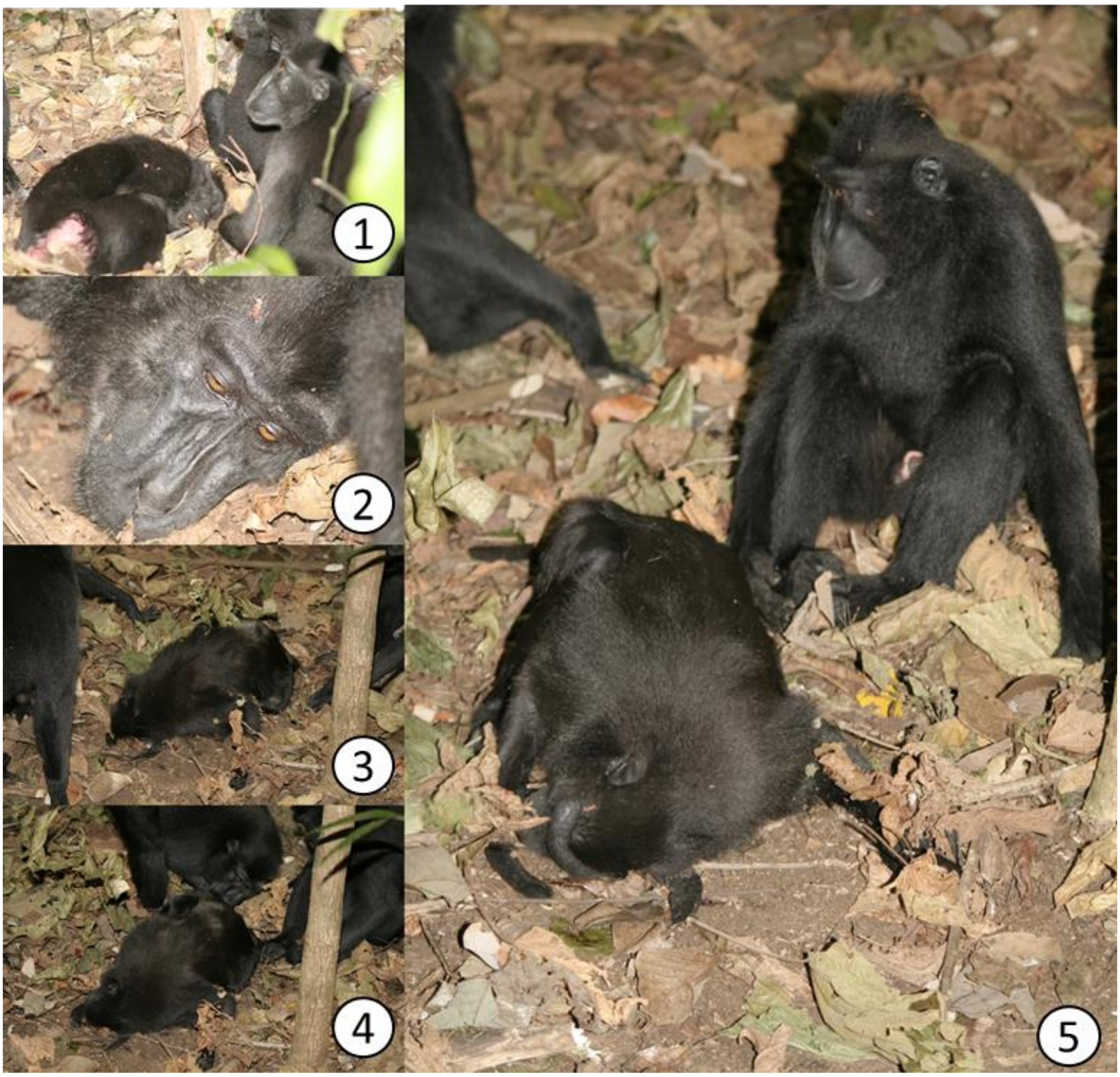

**Figure S1. Members of R1 attacking and adult female from R2 (Amber, AD) the 3rd August 2006. Pictures are numbered in chronological order.** Pictures by Antje Engelhardt. See descriptions of each image in the previous page

*Field notes*

The harassment lasted for about 45 minutes and was carried out by several juveniles, females, and sub-adult males. Amber survived.

#### <u>15th November 2006: R1 harassed an adult female from R2 (Betty, BD)</u>

**Available information about the attack:** 6 pictures of injuries, notes on the long-term demography database of MNP, two entries of intergroup encounters of the 15th November 2006 and personal observations by Antje Engelhardt.

*Pictures*

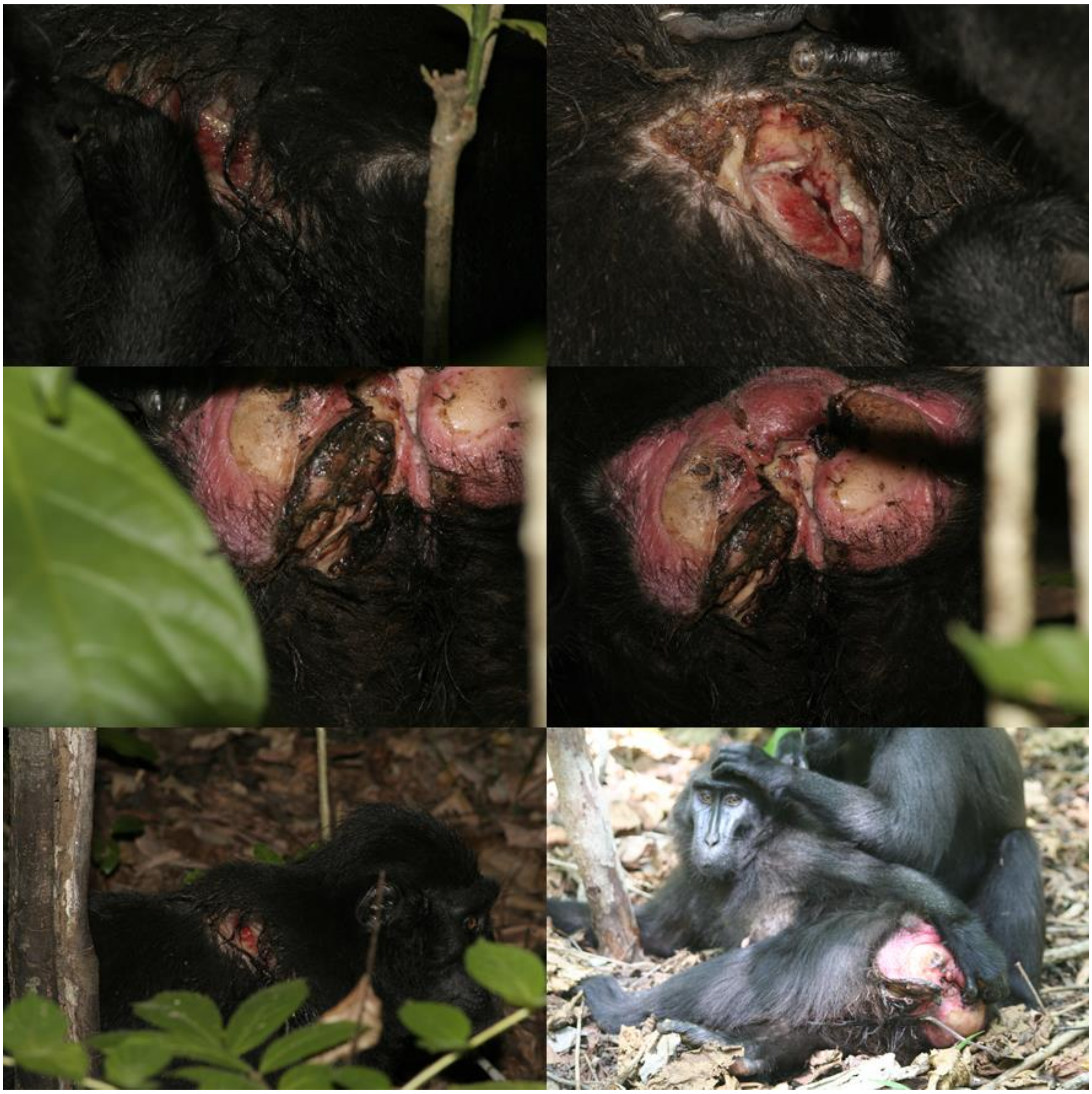

**Figure S2. Injuries displayed by an adult female from R2 (Betty, BD), after being attacked during intergroup encounters the 10th November 2006 and the 15th November 2006.** The pictures were taken the 15th November before Betty disappeared. Unknown photographer.

*Demography database entry*

*"10.11.06: severely wounded;15.11.06: harassed again. since then disappeared; around 20.11.06: infant disappeared (BD1B)"*

*Intergroup encounter entries*

**Table S2. Intergroup encounter entries of the 15th November 2006**

| Time: | Group 1 | Group 2 | Retreat: | Location: | Event: |
|---|---|---|---|---|---|
| **5:55** | RI | RII | RI | e 1000 | RII:c->RI:l |
| **10:15** | RI | RII | RI | aa-ab 5 | RII:c->RI:l |

*Personal observations*

*"The case of BD from R2, who end of 2006 went missing from the group straight after an intergroup encounter with R2. We found her a few days later dead."* Antje Engelhardt

#### 2007: R2 attacks an adult female from R1 (Bea, BS)

**Available information about the attack:** Personal observations by Antje Engelhardt

*Personal observations*
*"BS from R1 in 2007 getting trapped by adult females and adolescent males from R2. She finally managed to escape into the sea."* Antje Engelhardt

#### 18th July 2007: R1 harassed an adult female with infant from R2 (Indah, ID)

**Available information about the attack:** 10 pictures of the attack from an unknown author. An entry on the intergroup encounter record by Teija Febranouva.

*Pictures*

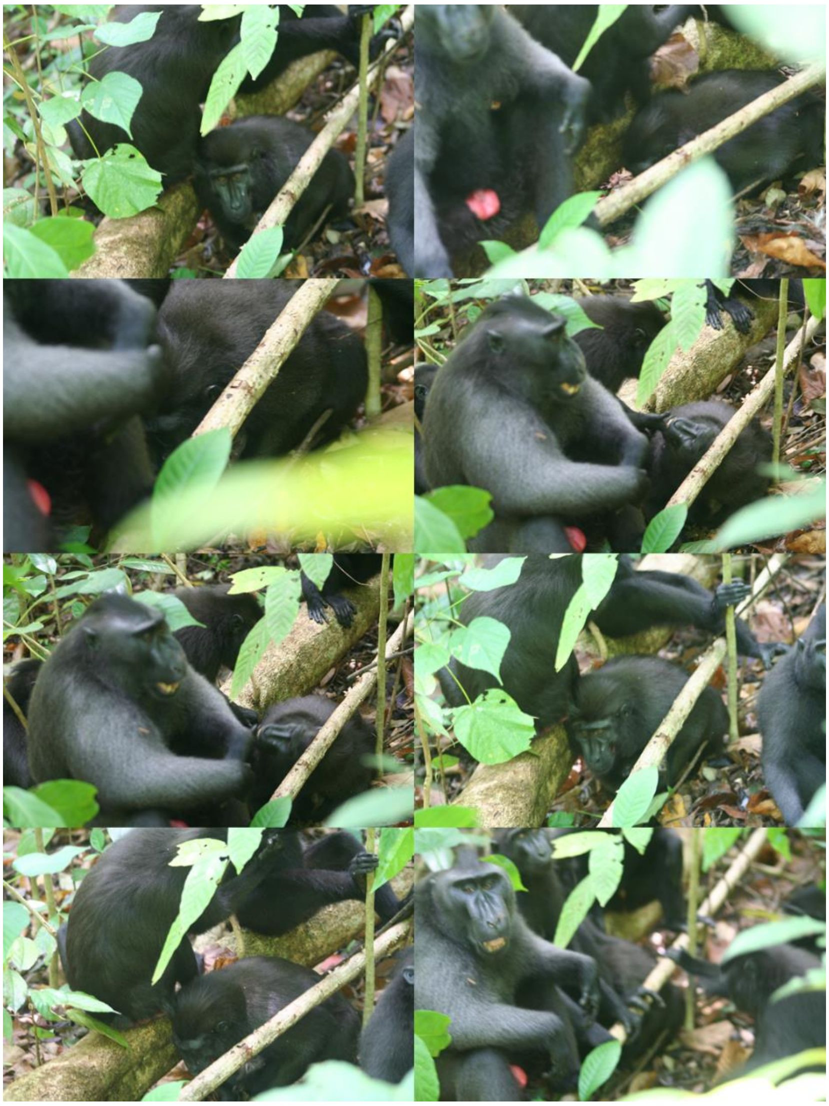

**Figure S3. Members of R1 members attacking and adult female from R2 (Indah, ID) and her infant.** The attack took place the 18th July 2007. The author of the pictures is unknown.

The pictures in **Figure S3**. show Indah curled up on the ground, apparently protecting her infant. She has a round wound on the left side of her mouth of a couple of centimetres of diameter. She is surrounded by several individuals, including an adult male, juveniles, and adult females. In two of the pictures, an individual is taking Indah's chin with its hand making her look at it.

*Intergroup encounter entry*

**Table S3. Intergroup encounter entries of the 18th July 2007**

| Time: | Group 1 | Group 2 | Retreat: | Location: | Event: |
|---|---|---|---|---|---|
| **6:39** | R1 | R2 | R1 | e1000 | |

#### 8th October 2007: R2 harassed an adult male from R1 (Bima, IJ)

**Available information about the attack:** 24 pictures from an unknown author taken during the attack. Sixteen of these pictures are attached in this document. Two entries in the long-term MNP database about individuals, an entry on the long-term MNP database on Demography by Gholib Assahad. There were no intergroup encounters recorded this day, which, together with the other entries, led to the inference that the victim had been travelling alone, which in not uncommon for males checking groups before migration.

*Entries in the database about individuals*

**Table S4. Entries in the database about individuals on Bima (IJ) regarding the harassment of the 8th October 2007**

| **ID** | IJ | IJ |
|---|---|---|
| **name** | Bima | Bima |
| **sex** | m | m |
| **group** | R1 | R1 |
| **date** | 08/10/2007 | 08/10/2007 |
| **event** | injury | missing/disappears from group |
| **detail 1** | | G200 |
| **detail 2** | | |
| **detail 3** | | |
| **detail 4** | | |
| **observers** | | |
| **notes** | | disappeared since 2.10.?; 8:00; near R2, harassed; IJ crouch and sbt.(?), groomed by juvs |
| **latest change by** | siobhan | siobhan |
| **date of change** | before June 2014 | before June 2014 |

Sbt=silent-bared teeth

*Pictures*

There was a total of 24 pictures available. Three were discarded because they only showed the unwounded face of the victim. Five were discarded because they were taken within seconds of those displayed in **Figure S4.** and **Figure S5.** and did not add information.

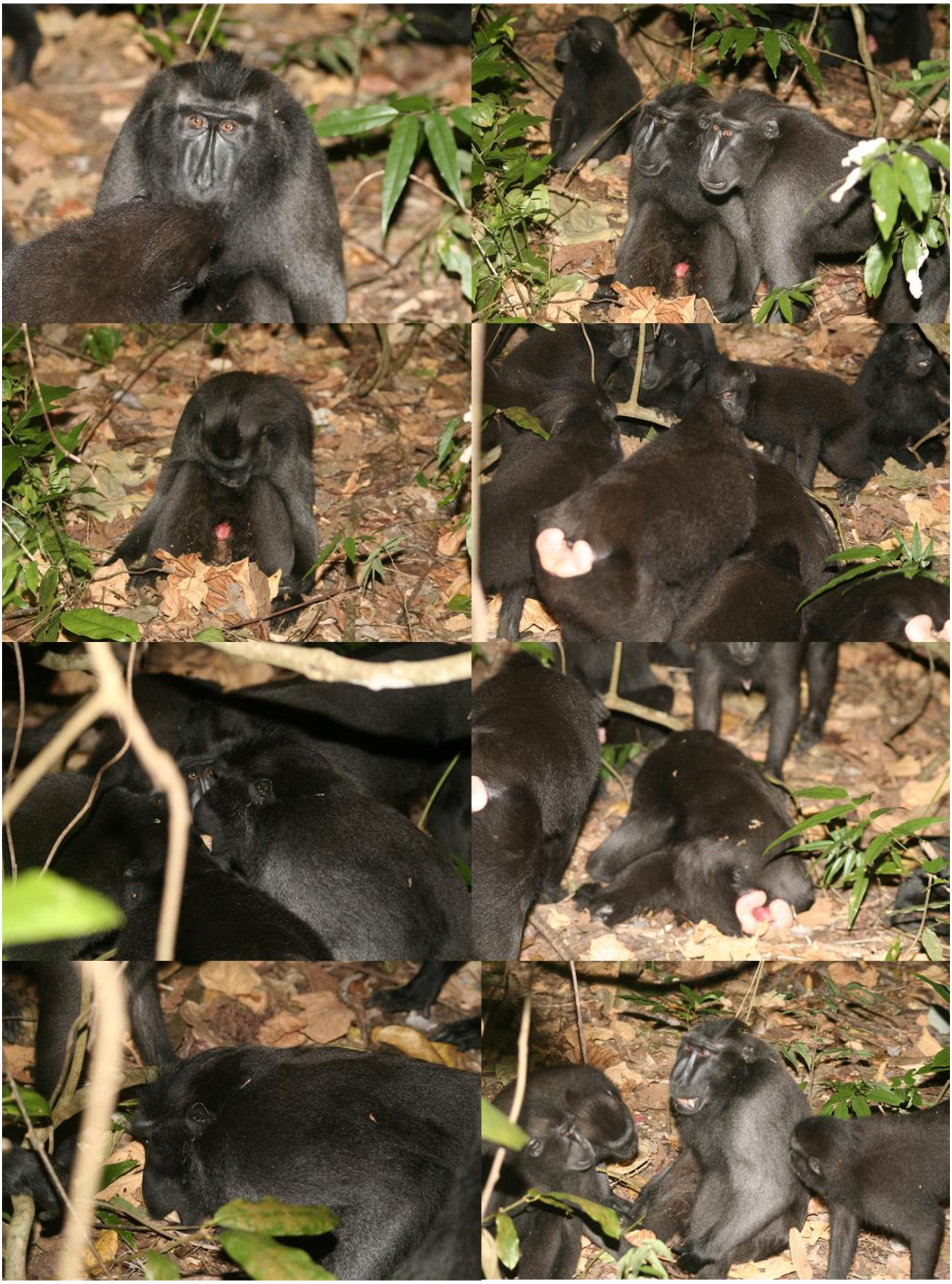

**Figure S4. Members of R2 harassing an adult male from R1 (Bima, IJ) the 8th October 2007.** Unknown author.

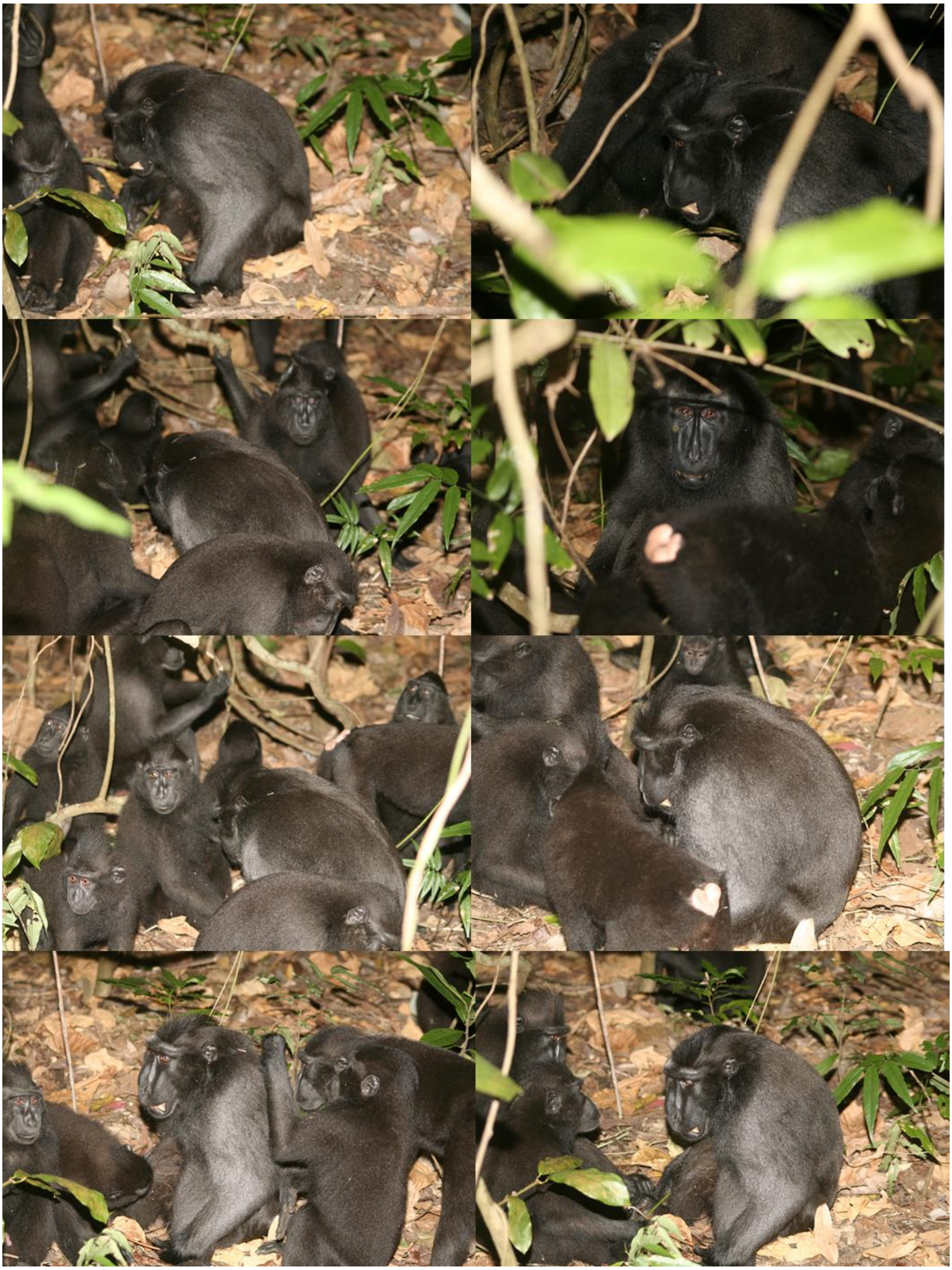

**Figure S5. Pictures taken while R2 harassed and adult male from R1 (Bima, IJ) the 8th October 2007. Unknown author.**

The pictures in **Figure S4.** and **Figure S5.** show the adult male from R2, Bima (IJ) usually in the centre of the scene. He is either sitting or crouching on the ground, often with silent bared teeth. Around him, there are mostly juveniles as well as at least one adult female (based on elongated nipples) and one adult male (based on similar size to that of the victim). All of them are inferred to be from R1 as for the accompanying entries (see below). No attacks are seen in the pictures. Instead, we can see juveniles grooming the victim.

*Demography database entry*
*"08.10.07 around 8:00 G200: R2 meets IJ, R2 harassed IJ, IJ crouched and sometimes silent-bared teeth. Some juveniles grooming IJ. IJ limping after"*

#### 14th November 2007: Adult female with infant of unhabituated group attacked by females from R2

**Available information about the attack:** A note on the intergroup encounter long-term database by Teija Febranouva

*Intergroup encounter entry*

**Table S5. Intergroup encounter entry of the 14th November 2007**

| Time: | Group 1 | Group 2 | Retreat: | Location: | Event: |
|---|---|---|---|---|---|
| **9:35** | R2 | Kalibersih | Kalibersih | aa6 | SJ in Kalibersih group, one female from Kalibersih was being harassed by R2 female, so many wound, UJ wounded on his left leg, HD wounded on the face and swelling and MD infant wounded on the back |

#### 14th November 2007: R1 attacks adult female (Maria, MD) with infant (MDB1) from R2

**Available information about the attack:** An entry in the long-term MNP demography database under the individual Maria (MD), as observed by Teija Febranouva (TF), Julianus Mendome (JM), Gholib Assahad (GA), and Antri (AN). An entry in the long-term MNP intergroup encounter database by Teija Febranouva. Pictures of the infant MDB1 after being wounded, presumably during a previous encounter with an unhabituated group (Kalibersih, see information on the previous attack).

*Pictures*

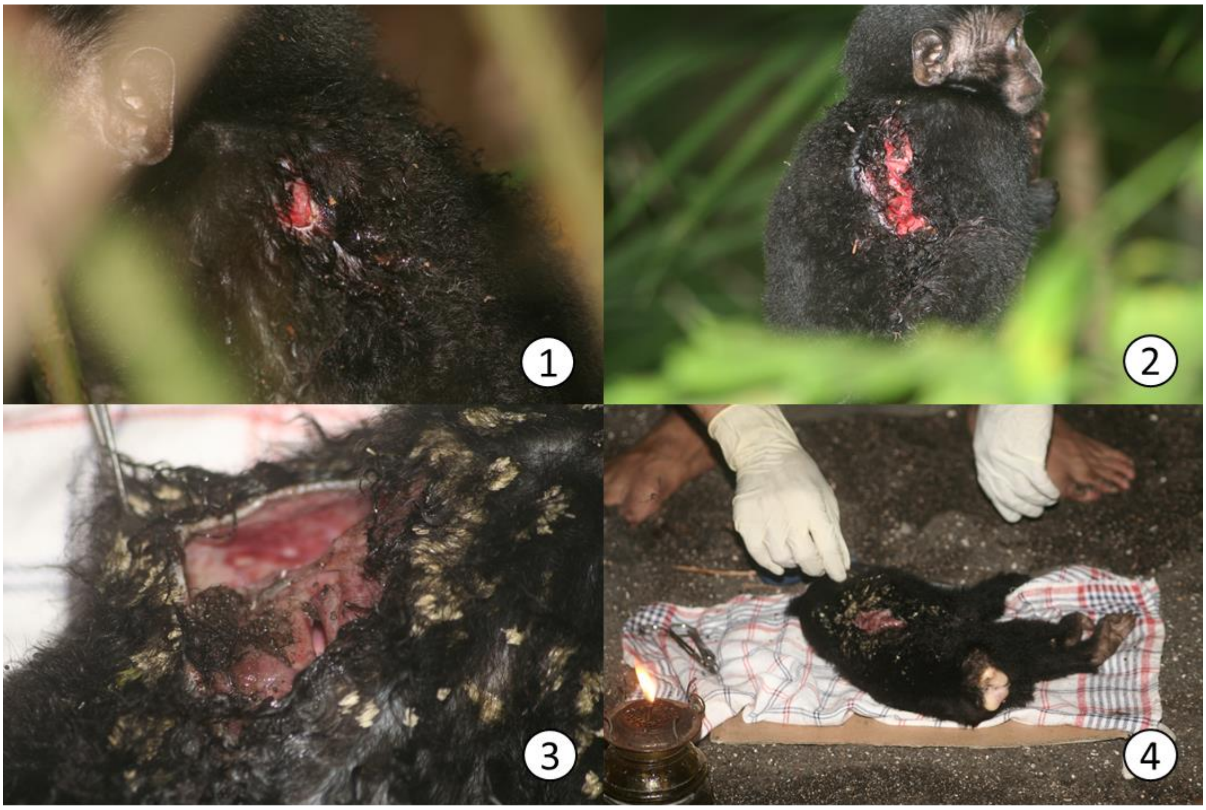

**Figure S6. Pictures of the injuries of the infant from R2 (MD1B) after an encounter with an unhabituated group.** Pictures 1-2 were taken the 14th November 2007 after the encounter with the unhabituated group. Pictures 3-4 were taken the 15th November 2007 once the infant died at the MNP camp POS3. Unknown author

*Intergroup encounter entry*

**Table S6. Intergroup encounter entry of the 14th November 2007**

| Time: | Group 1 | Group 2 | Retreat: | Location: | Event: |
|---|---|---|---|---|---|
| **11:30** | R2 | R1 | no | ab4 | |

#### Demography database entry

Under information for MD: *"14.11.07 : infant severely wounded on the back during intergroup encounter with Kalibersih group. left in R1, being harassed by QS, subadult, LJ and some juveniles, slept alone at AA4 while the group slept at AA2 (TF, JM); 15.11.07 Infant (MD1B) found in A600, with so many flies on the back around the wound. We tried to bring to POS 3 to save her (clean and give betadine), but suddenly she died (10 minute later) (GA, AN).”*

#### 16th February 2008: R1 harasses an adult female with an infant from an unhabituated group

**Available information about the attack:** Three non-consecutive video-clips of a total length of 2 minutes 54 seconds without sound by an unknown author. An entry in the long-term MNP database on intergroup encounters. The observer was Teija Febranouva

*Videos*
Three videos compiled in **Video S1**. Here is a narration of them:

The first clip (26.5 seconds): A sub-adult male and what looks like three juveniles and an adult female from R1 harass a female with an infant from an unhabituated group. The sub-adult bites her on the back, and the female from R1 grabs her. Then the victim flees twice for a few meters, but the aggressors quickly catch her both times. When grabbing her the second time, a sub-adult male jumps on her back and runs away, while the female and another individual bite the victim. When the female aggressor turns around, she has fur on her mouth. A juvenile grabs some of the unhabituated female's fur from the back. Then the victim runs away again, showing the long cut (~10 cm) that she has on her left flank.

She is caught again by a juvenile and a sub-adult, who presses her against the ground. A juvenile smells her and then grabs her fur and pulls and tries to examine her face, but the sub-adult grabs the victim and pulls her toward him. The sub-adult bites her and another one together with a juvenile examine her. One of the sub-adults grabs fur of her back and pulls, and in doing this, the victim tries to run away again. She manages to not being caught up to the end of the video.

The second clip (1 minute, 19 seconds): The victim is curled up on the ground and a female and a sub-adult male manipulate her body, while a juvenile watches. The sub-adult leaves. Then the juvenile and the female bite the victim. The victim tries to run away, but the aggressors trap her. Two juveniles bite her, and a third individual touches her back. The juveniles bite her again. Then the female and a juvenile, while a third individual wraps its arm around her. The female bites the victim's arm. Then the sub-adult and a juvenile seem to bite as well. More individuals join the attack and soon is not possible to see who is doing what or even how many they are precisely, but at least seven.

Suddenly, most of the aggressors flee, and the victim is left alone on the ground laying down. The sub-adult and a female are still close to her. The female moves away and the sub-adult male stays manipulating the victim, rolling her on the ground. The female comes back and grabs the victim's crest and pulls. The sub-adult drags the victim by pulling from one of her arms. Then he stands on her. When he steps down, the female approaches the victim with three juveniles. They start attacking the victim. They manage to turn her upside down, and when they do, they start biting her abdomen violently, particularly the sub-adult. She does not resist at any point. When the aggressors start leaving, the victim manages to lay facing down. Only a small juvenile stays with her; it puts its hand on her head and then jumps away. The victim then turns the head slightly, showing a sizeable wound on the right side of her forehead before she escapes.

The third clip (45 seconds): A crowd of juveniles, an adult female and a couple of sub-adult males hide the victim. Some individuals seem to bite her, but it is unclear. A female grabs the victim's back and then the victim defecates, while is attacked by juveniles. The female bites her as well. The victim moves slightly, trying to put all her limbs under her body. Then she is turned upside down by her aggressors, showing the long cut on her left flank. Two juveniles bite her, and a third individual grabs her crest and drags her towards itself. There is blood on her sexual skin. A couple of juveniles touch and smell her. A juvenile grabs her leg and drag her towards it and then turns her upside down. It and another juvenile bite her. She has blood on her face. She is dragged by a female and when this releases her, the victim flees and is chased by a juvenile and a sub-adult.

*Intergroup encounter entry*

**Table S7. Intergroup encounter entry of the 16th February 2008**

| Time: | Group 1 | Group 2 | Retreat: | Location: | Event: |
|---|---|---|---|---|---|
| **10:29** | R1 | kalibersih | Kalibersih | ab6 | 1 female with infant caught and being harassed by Ff and Jj R1. {BS, TS,QS HS etc}. the female escaped wt so many wounded while the infant lefted in R1. carried by R1 adolescent female. |

#### 16th June 2008: R1 harassing an infant from R2 (GD1B)

**Available information about the attack:** 2 entries on the long-term MNP database on individuals plus confirmation (email communication) with one of the observers (Julie Duboscq).

*Individual macaques database entry*

**Table S8. Entries in the database about individuals on GD and GD1B referring the attack the 16th June 2008 and its presumable consequences**

| **ID** | GD | GD1B |
|---|---|---|
| **name** | Ginger | -- |
| **sex** | f | male |
| **group** | R2 | R2 |
| **date** | 16/06/2008 | 21/06/2008 |
| **event** | infant injury | death |
| **detail 1** | GD1B | |
| **detail 2** | head | injuries to head |
| **detail 3** | intergroup encounter | |
| **detail 4** | R1, R2 | |
| **observers** | JD, ML | AN |
| **notes** | Pos3; harassed by juveniles, females; GD harassed by FJ | |

*Personal communication*

*"Ginger's (GD) infant (male), from R2, was isolated during an intergroup encounter between R1 and R2. It was harassed by several females and juveniles. His mother, GD, was chased by an adult male (Barra, FJ). Both, mother an infant returned to their group, but the infant was severely wounded on his head and died 5 days after the attack.*" Julie Dubosq

#### 2nd November 2008: R1 harassing a juvenile of either R2 or R3 (triadic encounter)

**Available information about the attack:** 4 pictures and field notes by Jerome Micheletta

*Pictures*

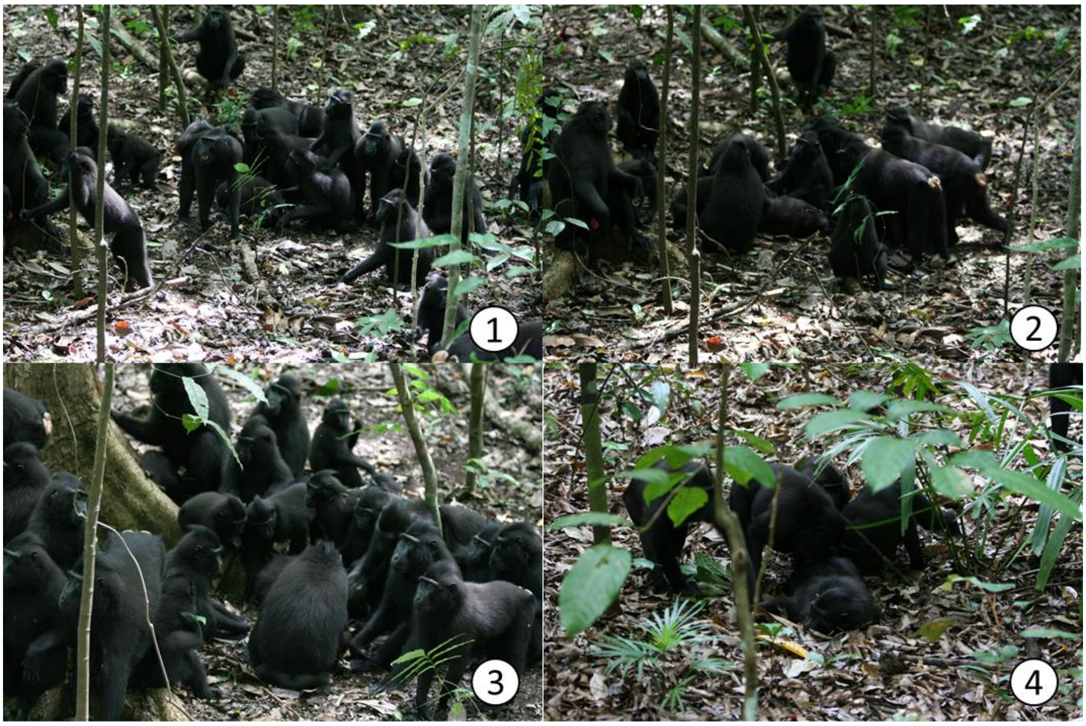

**Figure S7. R1 harassing a juvenile from either R2 or R3 the 2nd November 2008.** Pictures by Jerome Micheletta

In **Figure S7.1** and **Figure S7.2**, we see the victim curled up on the ground surrounded by juveniles, sub-adult, adult females, and a single adult male which is within 2 meters of the victim. Some individuals touch the victim. In **Figure S7.3** the victim is hidden behind a female from who we only see the back. The victim is surrounded by juveniles and at least one subadult male and two adult females. In **Figure S7.4** we see the victim lying down on their stomach, surrounded by 4 immature individuals examining them.

*Field notes*

*"1j R2/3? harassed bleeding when escaped"* Jerome Micheletta

#### 1st December 2008: R1 attacks a juvenile from R2

**Available information about the attack:** Field notes by Jerome Micheletta.

*Field notes*

**Table S9. Field notes on the attack of the 1st December 2008. They were recorded by Jerome Micheletta.**

| Note | Interpretation |
|---|---|
| *"bs es ns qs xj ar jR2"* | Bea, Erna, Nuria, Quty and several juveniles harass a juvenile from R2<br>(Bea, Erna, Nuria, Quty were adult females from R1) |
| *"jR2 escape after 15mn saw no injury"* | The Juvenile from R2 escaped after 15 minutes. No injury observed. |

#### 20th December 2008: Juvenile PB attacked by R1

**Available information about the attack:** Field notes by Jerome Micheletta

*Field notes*

**Table S10. Field notes on the attack of the 20th December 2008. Recorded by Jerome Micheletta.**

| **Note** | Interpretation |
|---|---|
| *"lsbsnsxjarjPB"* | Leoni, Bea, Nuria and several juveniles harass a juvenile from PB<br><br>(Bea, Erna, Nuria, Quty were adult females from R1) |

#### 5th February 2009: R1 attacks adult female from PB (Agnes, AP)

**Available information about the attack:** Field notes by Jerome Micheletta

*Field notes*

**Table S11. Field notes on the attack of the 5th February 2009. Recorded by Jerome Micheletta.**

| Note | Interpretation |
|---|---|
| *"just saw qs xj sa bs xs ar&ab ap"* | The observer only saw Quty, several juveniles, a subadult male, Bea and BigNose harassing and biting Agnes (Quty, Bea and BigNose were adult females from R1 and Agnes an adult female from PB) |

#### 31st May 2009: R1 attacks an Adult female from R3

**Available information about the attack:** Field note by Jerome Micheletta

*Field notes*

**Table S12. Field notes on the attack of the 31st May 2009. They were recorded by Jerome Micheletta.**

| Note | Interpretation |
|---|---|
| *"xfxsaarf3, start in trees"* | Several adult females and subadults harass an adult female from R3. The attack started on the canopy. |
| *"xf: qs.ns.zs.bs"* | The attacking adult females were: Quty, Nuria, Zoe and Bea. |

#### 24$^{th}$ November 2009 Juvenile R1 attacked by PB

**Available information about the attack:** Field note by Dwi Yandhi Febriyanti

*Field notes*

**Table S13. Field notes on the attack of the 24$^{th}$ November 2009. Recorded by Dwi Yandhi Febriyanti**

| Note | Interpretation |
|---|---|
| "xjarj" | Several juveniles harass juvenile |
| "xj=pb" | The attacking juveniles belonged to pb |
| "j=r1" | The victim juvenile belonged to r1 |

#### 1st January 2010: R1 attacks adult male from R2 (Vlad/Victor jr, VL)

**Available information about the attack:** Note in the long-term MNP database on demography and entries in the long-term MNP database on individuals.

*Demography database entry*

*"02.01.10: not in R2 (harassed by R1 the day before? IR); 03.01.10: VL wounded on the back & right leg, left arm limping (from IE R1 on the 1st?). chased away from R2 by males, some females and juveniles (NC, DZ)."*

Observers: Ira Ratna Sari (IR), Nicole Aline Thompson (NC), Deidy Azhari (DZ)

*Individual macaques database entry*

**Table S14. Entries in the database about individuals on Vlad/Victor Jr (VL) referring the attack the 1st January 2010 and its presumable consequences**

| **ID** | VL | VL | VL | VL |
|---|---|---|---|---|
| **name** | Vlad (Victor Jr) | Vlad (Victor Jr) | Vlad (Victor Jr) | Vlad (Victor Jr) |
| **sex** | m | m | m | m |
| **group** | R2 | R2 | R2 | R2 |
| **date** | 02/01/2010 | 02/01/2010 | 03/01/2010 | 03/01/2010 |
| **event** | missing/disappears from group | back to group/reappearing | injury | missing/disappears from group |
| **detail 1** | | | back, right leg, | left arm |
| **detail 2** | attacked/harrassed | | intergroup encounter | chased |
| **detail 3** | | | R1, R2 | by females, juvs |
| **detail 4** | | | | |
| **observers** | IR | | | NC, DZ |
| **notes** | by R1 on Jan 1 | | on Jan 1 | |
| **latest change by** | siobhan | siobhan | siobhan | siobhan |
| **date of change** | before June 2014 | before June 2014 | before June 2014 | before June 2014 |

#### September 2011: Juvenile R3 attacked by R1

**Available information about the attack:** Notes in long-term MNP database on demography

*Demography database entry*
*"Juvenile R3-Small juvenile was harassed in R1 for more than 3 hours".*

Observers: Meldy Tamengge (ML), Benediktus Giyarto (Ugik, UG), Julianus Mendome (JM).

#### 18th December 2015: PB1 lethally attacks and outgroup female

**Available information about the attack:** Three consecutive videos of the attack. They have a total duration of 35 minutes and 30 seconds and recorded by Laura Martínez-Íñigo. Fragments have been edited together in **Video S2**. A summary of the event and a full narration of the video can be see below as in the appendix of *Martínez-Íñigo, L (2018). Intergroup interactions in crested macaques: factors affecting intergroup encounter outcome and intensity". Ph.D. Dissertation, University of Lincoln*

*Summary*

**Observer/s:** Laura Martínez-Íñigo (LI), Rismayanti (RY).

**Victim:** Adult female of an unhabituated group

**Aggressor group:** PB1

I (LI) was following PB1 with four other observers (Rismayanti, (RY), Meldy Tamengge (ML), Meydi Ardi Manderos (MS) and Iwan Halir (IW)), and I was looking for my next focal. The group was very spread out, and I lost contact with the monkeys for some minutes. When I finally found some of them again (10:48), they were attacking a female from an unhabituated group. A juvenile male[1], called Sashimi, and an adolescent female, Emping, were the main attackers. They both bit the victim repeatedly through the whole observation period. Some unidentified juveniles also bit the victim, whenever the other two macaques allowed it. An adult female, Jane, was seen biting the unknown female twice. While Emping and the juveniles focused their bites on the female's limbs, Sashimi did so on her throat. Possibly, one of those bites to the throat killed the female by suffocating her. There was one in particular, at 10:54:45, that lasted for 15 seconds, that seems the most likely candidate. Before this bite, the female did not struggle and played death most of the time, although it was possible to see some limb movement every now and then. The most apparent movement occurred at 10:52:50, when she turned herself upside down. After the bite mentioned above, I saw no more movements, although I could detect a shallow breath which is not remarkable in the videos.

[1] Sashimi was almost a sub-adult. He was bigger than a female but he still lacked other features to categorize him as sub-adult. Those were elongated canines and reddish skin on the anal area.

Besides bites, some attacks consisted of pulling parts of the victim's body and dragging her. These were a minority in comparison to the bites and occurred mostly after the presumed death.

Many bystanders examined the body, sometimes touching briefly and/or sniffing. There were periods in which nobody interacted with the victim and the monkeys around continued normal activities.

There were two moments in which the victim was left alone. The first one occurred at 10:59, when the victim was left for 1 minute and a half, after Kristi, one of the adult females had been examining her. Interestingly, she came back vocalizing softly, followed by Sashimi, who bit the victim on the abdomen. Meanwhile, Kristi vocalized excited, and Emping appeared running and joined the new bout of attacks, together with a couple of juveniles. Around 11:15, the victim was again left on her own for a bit more than a minute. Then Sashimi and Emping returned for a while and left again. Finally, Sashimi came into visual contact again with the corpse but left shortly afterwards not to return.

The examination of the body showed superficial injuries on her right arm (2-3cm), two on the groin (~3cm each), several lacerations of about 1cm each on the chest and one on the throat of ~1.5 cm (see **Figure S9**). None of them bled and seemed only to pierce the skin. If the cause of death were the attack, I would presume it was due to suffocation caused by the bites directed to her throat by Sashimi.

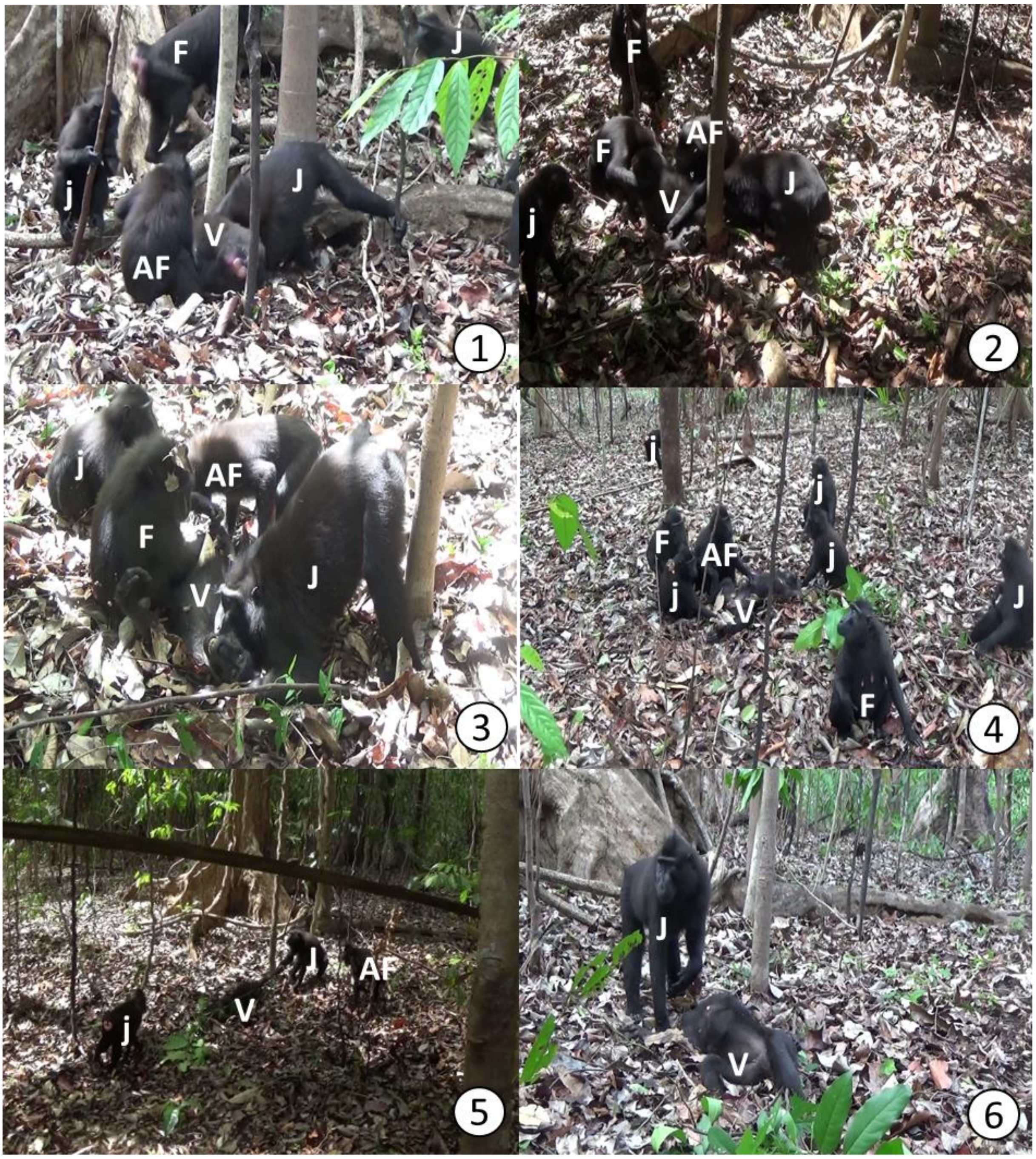

**Figure S8. Members from PB1 attacking a female from an unhabituated group on the 18th December 2015.** Pictures numbered in chronological order. Age sex classes are labelled as: AF= Sub-adult female (In this case always Emping); F= Adult Female (In this case always Jane and Bianca); j=Juveniles J=Juvenile (Sashimi); V=Victim. Pictures by Laura Martínez Íñigo

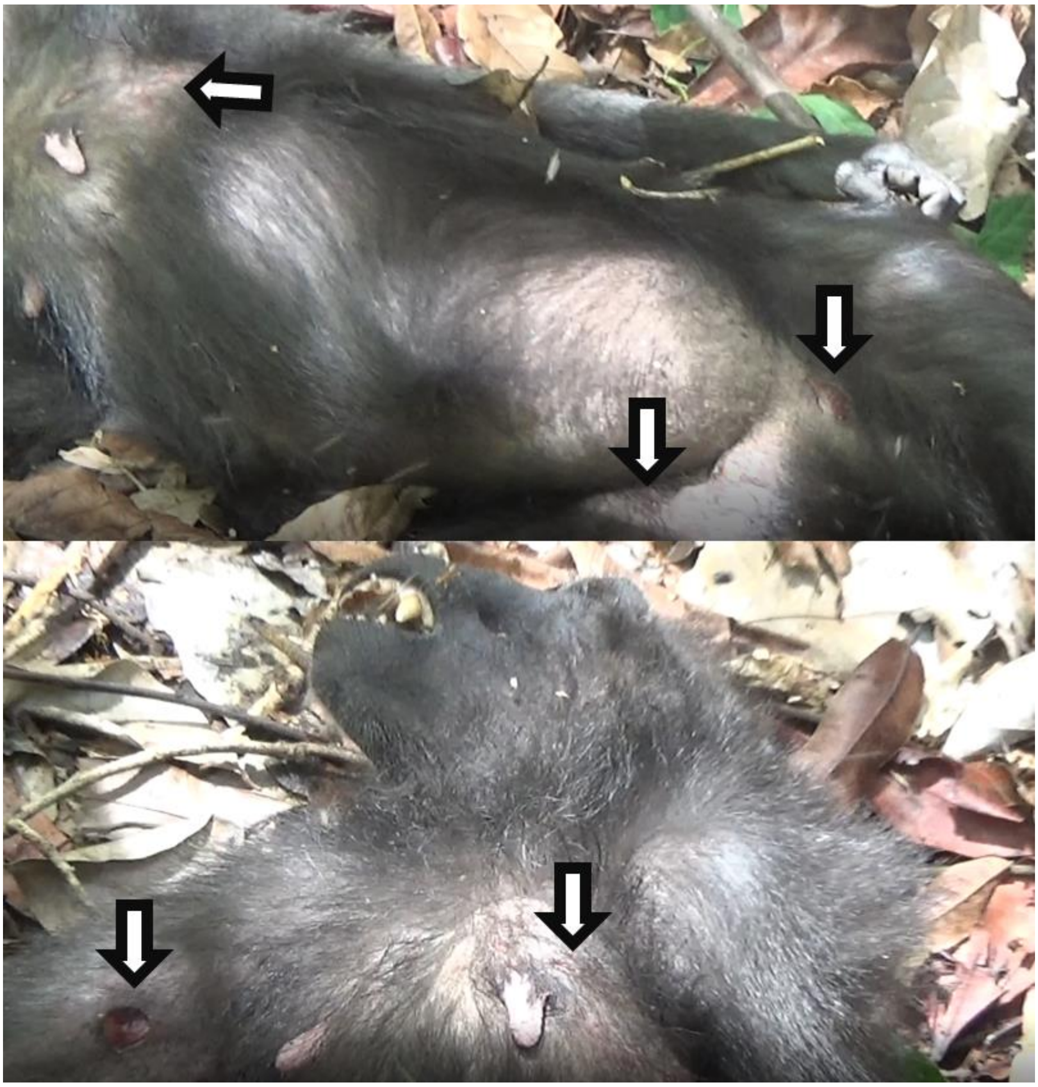

**Figure S9. The corpse of the unhabituated female attacked the 18th December 2015.** The arrows point at wounds. Pictures by Rismayanti.

*Full description*

It was the 18th December 2015 at 10:48. I (LI) was following PB1 with 4 other observers (RY, ML, MS, IW), and I was looking for my next focal. The monkeys were very spread out, and I lost contact with any of them for some time. When I finally found some more monkeys, they were already attacking the female (UF=Unknown Female), and I had started recording the event by 10:49.

At that point, within 2 meters of the victim, there were 2 juveniles, 2 individuals that could not be identified, 2 adult females (Jane, JP, and Bianca, BP), the sub-adult female Emping

(EA, AF in **Figure S8**) and the juvenile Sashimi (J in **Figure S8**). BLS=Body lengths (average female body-length from shoulders to the beginning of the tail).

Sashimi and Emping bite the female violently. The female does not struggle but moves her hind legs slightly. Meanwhile, Bianca displaces juveniles around the victim, while Sashimi and Emping keep biting the victim at intervals (**Figure S8.1**). A juvenile approaches but is displaced by Bianca. Emping hugs Bianca, and she reciprocates. By now, only Bianca, Sashimi, Emping, Stephie and a juvenile are within 2 m of the female. A minute later, Sashimi leaves and two juveniles touch the UF, pulling her arms and nipples. She does not respond. Sashimi grabs the left arm of the victim and pulls towards him. The UF faces upwards with open eyes and grinning, looking dead. Emping bites the UF. Sashimi leaves and once the UF stops from being attacked, she turns around, lying on her stomach. Then, Sashimi appears and bites the victim on the throat, while Emping bites different parts at intervals.

Sashimi, who had left the scene, gets closer to the UF, bends and bites her left flank firmly while holding her head with his left arm, turns her around and steps on her head with his left foot. A juvenile and Emping pull from the female in different directions while Sashimi attacks her. Jane approaches, smells the female and sits. Emping moves towards the UF's face and then bites her and pulls her hair while Sashimi keeps biting as well.

Emping bites UF's left leg, Sashimi touches UF's head. And UF seems to try to stand up, moving her hind legs. Emping pushes the UF down with her left arm while still biting her. Jane joins, and so does Sashimi, grabbing UF's head (**Figure S8.2**). When they stop, Sashimi turns the female around, so the ventral part faces upwards. Then Sashimi bites the UF again between the chest and the throat. After, he sits down while UF faces upwards with open eyes and a grin. We see a juvenile, Jane, Bianca, Emping and Sashimi surrounding her. Then Sashimi grabs UF's lips and pulls them open before bending over UF's mouth to smell it. Jane and Bianca leave. Emping grooms UF slightly, then she bites her.

Sashimi grabs UF's right arm and pulls. Then he bends towards her and, while putting his left hand over her eyes, bites UF on the back of her neck followed by the throat (**Figure S8.3**). Emping and a juvenile are touching UF meanwhile. Sashimi stops biting UF's throat. The juvenile bites UF and Emping is in body contact with her. Another juvenile grabs UF's left arm and pull from her, displacing her. Then it leaves.

Emping bites again and Jane and Bianca appear in the scene. Two juveniles are seen as bystanders too. Emping drags UF while biting her. Sashimi and a juvenile follow through. Emping stops biting UF. A juvenile and Sashimi do as well. Bianca displaces a juvenile. A juvenile and Sashimi bite UF again, with Emping within one BLS, but obscured by Sashimi. Another juvenile approaches UF, grabs her head, pulls softly and grooms her. The other juvenile bites around UF's groin. Sashimi grabs UF's left leg, pulls and smells the foot.

A juvenile pulls UF towards itself, grabbing a leg and then dragging her. Another juvenile jumps around and a third one approaches to UF's head (not seen what it does). One of the juveniles bites the right flank and is then displaced by Emping, who ignores it, lip-smacks and approaches UF. UF lies facing upwards with a grin.

Emping bends and smells UF's mouth. A juvenile touches UF's right thigh. Emping displaces it and bites that area. Then a juvenile passes its hands over the area. Emping pulls UF's right nipple. Bianca and Jane are in the scene (**Figure S8.4**). Bianca is grooming the juvenile that grooms UF. The juvenile stops grooming UF. Jane leaves the scene. Bianca stops grooming a juvenile. Emping touches UF with both hands. A juvenile grooms UF.  A juvenile turns UF around. Another juvenile touches UF, and Emping examines UF's groin. UF is left resting on her right flank.

Sashimi bends and examines UF's head. Then he bites the throat firmly. Bianca leaves the scene and a juvenile grooms UF's hindquarters. Sashimi leaves the scene running, and a juvenile follows. Then Emping moves away from UF. A juvenile then bends towards UF's hindquarters and examines

We hear vocalizations typical of a conflict and Emping runs away from the scene. The two juveniles close to Emping look toward the same direction as Emping did. There is another individual over 5BLS away, on the background.

The juvenile who was in body contact with UF runs away from the scene. The juvenile who was on the background comes closer (2-3BLS). Then such a juvenile and another who was close to UF, run away from the scene. No monkey is seen around UF at this point. UF lies on her right flank; she seems to breathe. A juvenile appears in the scene and approaches. Looks around and bends to smell UF. The juvenile seems to touch UF (but then the camera moves and I cannot see). Then "smells" again and looks around. The juvenile touches UF, smells and moves around. The juvenile lip-smacks to someone out of scene and moves away from UF (1-2 BLS). Kristi appears and approaches UF.

Kristi bends and seems to smell UF, but her body covers what she does. The juvenile approaches Kristi teeth-chattering. Kristi lips-smacks at the juvenile. Then Kristi continues examining UF, the juvenile sits and grooms UF briefly.

Kristi smells the head of UF. The juvenile smells the left foot of UF. Kristi is sitting close to UF's head. Juvenile greets someone; I don't know if to Kristi or someone out of the scene. The juvenile grooms UF's hindquarters for a couple of seconds. The juvenile lip-smacks at Kristi and Kristi lip-smacks back. Then the juvenile approaches Kristi and smells UF head area. The juvenile grooms UF and Kristi stands up and leaves the scene. The juvenile follows. No more monkeys are visible in the proximity to UF.

Kristi approaches from over 5 meters away and vocalizes. A juvenile and Sashimi also approach. I retreat, moving away from UF. Sashimi approaches UF and bites her belly. Kristi approaches vocalizing. Emping comes close to UF, Sashimi and Kristi running. Sashimi smells UF's hindquarters, and Emping examines UF's head and bites her. Another juvenile next to UF examines her and seems to bite her abdomen. A second juvenile approaches, displacing the first juvenile. Sashimi bites the area of the head of UF, obscuring Emping. Both Emping and Sashimi bite around the head, while the juvenile that just approached smells the anal area of UF. Another juvenile approaches and touches UF's right arm and then bites. Another juvenile appears in the scene.

The juvenile who just appeared in the scene gets close to UF. Sashimi is sitting on UF's body. Emping moves away (but still within 0-1BLS) and Sashimi bites UF again around the head-neck area. Emping bites UF around her left arm. A juvenile seems to bite UF around the head too, but Sashimi and Emping are in front, so is not clear. Emping stands up while biting UF, pulling her arm up with her mouth, while pushing the body against the ground with her right arm. The juvenile that was close to Emping jumps away when she looks at it. Emping smells UF's anal area and sits in body contact. Sashimi stops biting UF. The juvenile that appeared biting before is no longer close to UF and Kristi is no longer in the scene. Sashimi leaves UF, Kristi is seen back in the scene vocalizing towards UF, Emping stands up on her hind legs and looks away, towards the direction Sashimi is heading. Then she runs that way, and Kristi looks towards that direction. Then Kristi looks at UF and sits within 2BLS of her. Two juveniles that were within 3BLS of UF leave, only Kristi remains visible in proximity. Sashimi approaches again, and Kristi vocalizes. Sashimi bends and bites UF around the left armpit. Sashimi stops biting and smells the anal area. Kristi keeps vocalizing. Emping comes back and touches UF's head, then grabs her face and pulls her head briefly upwards. A juvenile approaches and sits 1-2 BLS away of UF. Emping leaves. Kristi leaves. A juvenile approaches UF. Sashimi is in body contact with her. Then he stands up in two legs and looks away before sitting again. The juvenile smells UF. Sashimi moves towards UF displacing the juvenile. Then he bites the left flank of UF while the juvenile vocalizes. Afterwards, Sashimi tries to turn UF around (she is facing the ground at this point).

Sashimi puts UF's body on the left flank and grooms the abdomen for a moment. Sashimi runs away and disappears from the scene. Then the juvenile starts leaving, stopping briefly to observe UF.

A juvenile approaches UF, looks at her and grooms UF's abdomen briefly. Then the juvenile turns UF's body around and grooms the back briefly. A loud call is heard. The juvenile touches UF slightly, and another juvenile appears within 3BLS. Emping runs towards UF displacing the two juveniles that were close to her. She stops almost in body contact with her. Then Sashimi approaches too. Emping seems to bite slightly on the lower abdomen (not clear). Another juvenile approaches.

Emping bites UF abdomen and then moves away. Sashimi starts following Emping while grabbing UF's left arm and dragging her with him towards Emping (**Figure S8.5**). Then Sashimi, Emping and a juvenile examine UF, smelling and touching. Emping bites UF's right arm. Sashimi pulls the head up, looks the face closely and seems to lip-smack at her before biting UF's throat. The juvenile jumps away and Emping bites the lower abdomen. The juvenile approaches again, grabs UF's left leg and pulls towards him while Sashimi and Emping bite UF. A juvenile comes running towards them. The juvenile that was already there is also biting now. Emping leaves.

Sashimi grabs the right arm of UF and pulls from it. Then he moves away and stays within 3BLS. A juvenile smells UF's breast. The other juvenile approaches, smells and grooms slightly. One of the juveniles leave, the other grooms UF for a bit and then examines her anal area. The juvenile grooms UF again, and then grabs UF's left leg and smells her groin. Then the juvenile grooms UF's abdomen slightly. Another juvenile approaches and stays close for 20 seconds. I cannot perceive UF's breathing anymore. A juvenile grooms Sashimi within 5BLS of UF. A juvenile is around the UF's body, examining it. Then it bites it and grooms it. Then, after manipulating the body with its hands, sits, apparently in body contact with UF's corpse and waits. The juvenile grooms UF's body briefly again. Then the juvenile sits on the other side of the corpse. Vocalizations are heard, and the juvenile leaves UF.

Sashimi approaches UF again and bites her thorax and then her throat. The juvenile looks from 4-5BLS. Sashimi bites UF's right nipple repeatedly and pulls.  Sashimi turns the body around, examines the anal area and then pulls the body towards him, so UF's anal area is pointing upwards and her head and arms lie on the ground. Then he bites the groin. The juvenile approaches and after few seconds he bites the throat area while Sashimi bites tights and belly. Then the juvenile bites also the area between the abdomen and the tights. Sashimi bites UF's throat and the juvenile the groin area. Sashimi grabs UF's right arm and drags the body about 1BLS. Then he bites the throat, and the juvenile approaches the body again. There is another monkey about 3-4BLS away.  Sashimi pulls from one of UF's nipples. Then he stands up and "kicks" the body with his left leg. Then he moves away a couple of BLS. The monkey that was looking from 3-4BLS is a juvenile male. The juvenile close to the body bites the tights.  The juvenile moves away from the body (vocalizations are heard on the background). Sashimi also moves away from the scene. The other juvenile approaches the body and examines it before moving away.

Emping and Sashimi start approaching to UF again. Sashimi bites UF again on the abdomen-groin area. Emping is by his side, biting UF's left armpit area. Sashimi moves to bite UF's thorax and then he walks away. Emping follows Sashimi after smelling UF's body once more. They sit within 4-5 BLS of UF. There is a zoom into Sashimi's face, and when the camera zooms out at 6:14, Emping is gone. Sashimi stays within 5 BLS self-grooming and looking around.

Sashimi approaches UF body again and grooms her chest and shoulders (**Figure S8.6**). Sashimi leaves UF's body.

At the beginning of the third video, Sashimi approaches from far and looks towards the body from 5-10 BLS. Then he leaves and no more monkeys approach the body after this.

#### 20th February 2016: R2 female juvenile trapped by R1

**Available information about the attack:** Eleven videos of a total duration of approximately 1h15minutes. All videos were recorded by Laura Martínez Íñigo. Below, there is a summary of the event. Afterwards, there is a full description of what can be seen in the videos.

*Summary*

**Observer/s:** Laura Martínez Íñigo (LI), Juliette M. Berthier (JB), Julianus Mendome (JM), Meydi Ardi Manderos (MS), Santi Julianti (SJ), Try Sutrisno (TS)

**Victim:** Juvenile female from R2

**Aggressor group:** R1

R1 trapped a R2 female between the buttresses of a tree during an encounter between both groups. When the encounter finished, I (LI) started to cover the event. I could not observe in detail many of the aggressions that R1 directed to the juvenile from R2. The buttresses or the monkeys sitting on them usually covered the view obstructing the observation. However, it was apparent that at least Xiro (XK), two unidentified sub-adult females and several juveniles bit the victim. They manipulated the body of the victim, pulling her limbs and pushing her. The victim sustained the aggressions passively most of the time, playing dead. When the victim moved, it was to lip-smack at the monkeys around her and to turn herself down.

There was good visibility for two aggressions. In one of them, a juvenile male bit the upper part of the victim and pull her upwards until she was standing. Then he released her, and she dropped on the ground. In the other good-visibility aggression, a juvenile male bit the throat of the victim.

The day of the attack, we could individually recognize 48 individuals from R1. These were all the adults as well as some sub-adults. Of these 48 individuals, 25 approached the victim within one meter.  Five of these 25 were males. Tarzan (TM) and Aslan (AK) looked at the victim briefly and continued their activities. The other three males who closely approached the victim, Bento (BM), Ebleh (EJ), and Selatan (SN); looked at the victim for longer and stayed within proximity, but none on them went inside the tree gap to interact with the victim. Most of the adult females who approached the victim did the same as the adult males. The exceptions were Nihil (NU) and Adinda (AU). NU lip-smacked at the victim and went inside the gap, but I could not see whether she interacted there with the victim or not. NU groomed and embraced a juvenile from R1, who was in the gap within the buttresses close to the victim. At the time, the victim was curled between them and the trunk. AU went into the cavity and bent over the victim, but I could not see what she did there clearly.

It is worth noticing that on several occasions some sub-adult males of R2 were again within less than 100 meters of the closest members of R1.

In the end, the victim manages to escape while a juvenile was attacking her. A couple of R1 juveniles ran after her.  We could not find her again, and there were no reports of an wounded juvenile later that day in R2 nor the days after. We are uncertain whether she went back to her group or not. We were not able to see injuries during the observation.

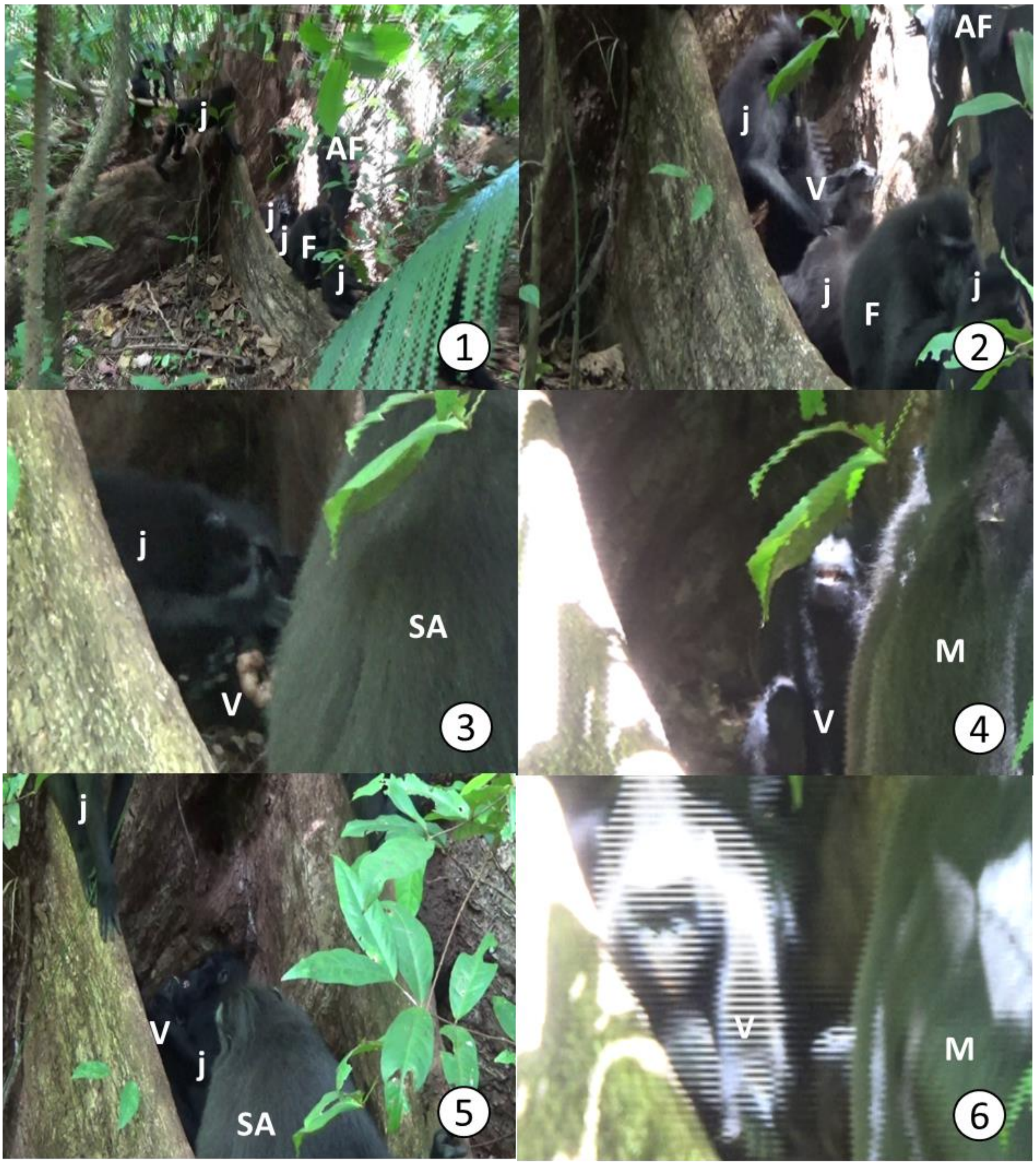

**Figure S10. Members of R1 attacking a juvenile from R2 the 20th February 2016, in chronological order.** Age sex classes are labelled as: AF= Sub-adult female; F= Adult Female; j=Juveniles; SA=Sub-Adult Male, M=Adult Male, V=Victim. Pictures by Laura Martínez Íñigo

*Full description*

It was the end of an encounter between R1 and R2. Me (LI) and JB were recording it when JM and MS let us know that a juvenile of R2 was trapped between the buttressed of a tree by R1. JB and I went there, and I started recording the event at 11:14. There were about 20 monkeys within 3-4 meters of the trapped juvenile of R2, including three females (Helena (HS), Nuria (NS), Gina (GU), Vodka (VS), Adinda (AU)), two sub-adult males (Niko (NK) and Xiro (XK)), two unidentified sub-adult females and many juveniles.  It took us time to find a proper place to observe the event, since the buttresses hid the juvenile and the individuals around, prevented us from approaching

At 11:17, I found a spot where I could observe a bit better and focus on the individuals seated on the buttressed around the victim (**Figure S10.1**). These were: NK, NS, GU and at least four juveniles. NK seems to look inside the buttresses and to lunge at the victim in a couple of occasions. NS leaves, and so does GU. I can only see NK and a few juveniles within 1m of the space between the buttresses. NK leaves, and a sub-adult female exists the space between the buttresses. She is quickly replaced by a juvenile who seems to bend to bite the victim. A sub-adult female displaces the juvenile. The victim is obscured by another adolescent and XK who arrive and put themselves in front of the scene. NK comes back, and another sub-adult female gets inside the gap between the buttresses. We hear screams in the video, but they come from behind the observer, not from the victim. XK leaves. A sub-adult female lip-smacks and presents to NK. NK examines her genitals but does not mount her. Instead, he presents her for grooming, and she does. After a while, Intan (IU), an adult female, approaches them and sits.  The sub-adult male Kevin (KK) approaches them displacing IU, who leaves. Then, there is a coalition against one of the monkeys. They redirect the aggression towards me, so I stayed still until they calmed down.

I started to record again at 11:27, but I could not find a spot to continue the observation until 11:30. A conversation with the observers following R2 revealed that the group was relatively close, enough as to be considered an encounter. Around the buttresses were the victim is, it is possible to see NK being groomed, but no more than this. I stopped recording and went to check where R2 was. At this point, JB had been less than 15 days in Tangkoko and was not ready yet to collect data, so she stayed watching whether the trapped R2 juvenile escaped or not while I checked the encounter. I did not see R2 close by, but I did see most of the group resting and grooming. They were spread out for several tens of meters, around the tree where the juvenile from R2 was. At 11:37, I see NK, SK and XK monitoring several sub-adults of R2 being. The rest of the R1 does not pay attention to those individuals from R2.

I see Kunti (KU) and Dadu (DU), adult females, approaching the area where the victim is. Then I check for R2, which is still 20-30 meters from the closest member of R1, who is Solo (SK), an adult male. Most of R1, however, is several tens of meters further away, mostly around the tree were the juvenile of R2 is being held. KK goes within the buttresses, but I cannot see whether he interacts with the juvenile from R2. There are several juveniles around the buttresses, and some vocalize, but the visibility is not good.

I find a better spot, and for a while, I only see KK's back while he sits on the tree buttress looking down at the R2 juvenile. By zooming, I see that someone else is with the juvenile in the cavity. I collect some data on the individuals around. When I point the camera again towards the buttresses, KK is gone. Instead, Lucifer (LK) is inside the cavity and manipulates the victim, who does not resist nor looks alive. LK goes out of the cavity. However, there is still someone else besides the victim. The unidentified monkey seems to manipulate the victim, but the visibility is low.

After a while, NU approaches through a third buttress. She looks down towards the juvenile of R2 and lip-smacks to her. Then NU leaves and Quty (QS), an adult female, passes by the buttresses. Aslan (AK), an adult male, also glances at the victim and leaves. Next, I go to check what the rest of R2 is doing. At 11:51, after looking around, I think the encounter is over and go back to where the trapped R2 juvenile is.

Iris (IB), a sub-adult female, goes inside the cavity, but I cannot see whether she does anything to the R2 juvenile. I get into a better spot, and I see a sub-adult female other than IB biting the R2 juvenile. A juvenile and AU are there, by the victim, as well.

After biting the victim, the sub-adult female grooms her briefly. LK approaches and goes inside the cavity, displacing the sub-adult female, who goes out and so does the juvenile. LK seems to bend towards the R2 juvenile, but only his back is visible. The sub-adult female that was attacking the victim leaves the proximity IB comes, and LK seems to be grooming the victim. IB leaves. A juvenile supplants IB and presents to AU, who grooms the juvenile.

A juvenile goes inside the gap while QS, who just approached, watches. Then, I did a group/scan before continuing with the recording of the events around the R2 juvenile. Bento (BM), an adult male, has come within proximity but leaves shortly after.  For a moment, I see a juvenile from R1 in the cavity lip-smacking to NU, who goes inside, embraces it, and grooms it. Then an infant goes into the gap as well. The juvenile from R2 is playing dead between the trunk and the juvenile from R1 that NU is grooming. When NU leaves, the juvenile from R1 grooms the victim, who stays still. Then, the juvenile grooms itself and Ola (OU), an adult female, come stands on the buttresses, preventing me for viewing the juvenile from R2. Afterwards, OU goes into the tree cavity. The juvenile that was already there keeps grooming the victim. A juvenile male mounts OU and gets inside the gap before mounting her again. Then both OU and the juvenile male leave the cavity. OU grooms the juvenile on the buttress in front of the cavity.

IB comes and looks inside the tree cavity. Another sub-adult female goes inside the gap, but I cannot see what happens there. A juvenile and the sub-adult female go out of the cavity. Next, a big juvenile male goes within the buttress roots and bites the victim. He bites the juvenile female from R2 on the upper part of the mouth, rising her until her body is straight (**Figure S10.2**). Then he releases her. Meanwhile, another juvenile in the cavity watches the attack. Both R1 juveniles go out from the tree cavity shortly after.

After a while, a sub-adult female and IB go into the cavity. One seems to be biting the R2 juvenile. The visibility for the other is bad, but she bends over the victim as well. Next, NU pulls IB's hair, and IB leaves the gap. The other sub-adult female stays, but soon OU moves, obstructing the observation. A juvenile descends then into the cavity, but I cannot see what happens inside.

When the juvenile leaves, I manage to get a view of the victim, who seems dead, lying down on one of her flanks. Then, it turns herself upside down and curls. A sub-adult female approaches the victim and touches her. She grooms the victim for a second, just before putting her foot on the victim's face. Then it is not very clear whether she bites around the mouth or only sniffs the area, but she manipulates the head of the R2 juvenile with her hands. Shortly after, a juvenile goes inside the tree cavity emitting play grunts and embraces the sub-adult female. Another juvenile joins them. The sub-adult female mock-bites the juvenile and then presents for grooming, which the juvenile does. The victim seems to be between them and the trunk, but it is not visible. NK approaches and goes inside the cavity but quickly leaves and goes out of sight. XK replaces NK and seems to manipulate the victim. Then XK goes out and sits on the buttress root in front. After a while, Polina (PS), an adult female, appears and presents to XK, who ignores her. Then Olivia (OS), an adult female, does the same and XK ignores her as well. OS leaves.

A sub-adult female looks towards the cavity while she is on a buttress root above. It looks like someone is biting the victim inside, but it is not possible to see it clearly with XK in front. Juveniles stand on the buttresses looking towards the victim. XK goes inside and seems to bite the victim. KU watches as well. Next, one of the male juveniles aggresses GU, who moves away and screams at him asking for support. Nobody comes, and she leaves. The juvenile male attacks another juvenile. The juvenile screams and asks for support, but the other macaques ignore him. The male juvenile who attacked GU and the previous juvenile tries to hit another juvenile and threats it. Meanwhile, he is covering what happens within the buttresses.

Cumi (CU), an adult female, passes by and stops a moment to look inside the cavity. Then she leaves. The male juvenile who was aggressing different subject moves away. Then we can see XK grooming the juvenile of R2, while a sub-adult female watches him doing so. XK stops and gets out, displacing another sub-adult female who was sitting on the buttress roots. He obscures the sight into the cavity. The sub-adult female leaves too. Shortly after, a juvenile male bites and rolls the victim. The juvenile from R2 does not struggle and plays dead. The juvenile from R1 keeps manipulating the victim's body, turning her upwards and pulling her legs up, biting and dragging her.

XK goes in, and then the victim lip-smacking at him. XK bends and seems to bite the victim, but the visibility is not good. XK goes out of the cavity and sits on the buttresses in front. The juvenile then starts biting the victim again, pulling her head upwards.

QS comes and presents to XK, who ignores her and presents her for grooming. She ignores him as well and leaves. The juvenile that was attacking the victim goes out of the cavity. Another juvenile pulls from the victim's hair, moving her whole body. Again, the victim does not resist at all and stays immobile facing down. A juvenile male gets inside the cavity, but quickly jumps out and sits on one of the buttresses. A couple of juveniles look inside the cavity standing. XK masturbates. Then he goes into the cavity and seems to do something to the victim, but I cannot see it. A sub-adult female approaches and sits on one of the buttresses.

AU and TM approach and look inside the cavity. TM loses interest almost immediately but AU peers inside for a while. QS approaches XK and presents. QS leaves and XK follows. AU sits where XK was, facing the cavity. The juvenile inside manipulates the victim's body, pushing and biting it (**Figure S10.3**). A couple of juveniles and a sub-adult female stay around the buttresses. Two juveniles mount the sub-adult female successively.

I notice that several individuals seem to be monitoring. I stop following the attack to the juvenile female from R2 to check whether anything else is happening. Soon, I go back after not finding anything unusual. When I continue the observation, Ebleh (EJ), an adult male, is sitting on the buttresses. He looks towards the cavity, where some individuals appear to be biting the victim. A couple of juveniles and a sub-adult female watch the aggression from the buttresses, and so does LK. Dori (DS), an adult female, passes by over the buttresses without paying attention to the attack. EJ leaves, and a juvenile goes into the cavity, where there is already a couple more. NU sits on the buttress roots facing the cavity. OU and a couple of sub-adult females are also on the buttresses over where the victim is. Nothing inside the cavity is visible. OU leaves, and so does one of the sub-adult females. AU goes into the cavity and bends over the victim but does not seem to attack her. Next, AU goes out and sits close. BM arrives and investigates the cavity. There are several juveniles inside around the victim, whom I cannot see. BM sits and grooms someone on the buttress root. A male juvenile goes out of the cavity. Super-nose (SN), an adult female, climbs the buttress and looks inside the cavity; then leaves.

Selatan (SN) stands over the buttresses and peers inside the cavity as well. Next, SN approaches BM, displaces him and supplants him. OS approaches and presents to SN, who examines her and starts grooming her. A juvenile seems to be in contact with the victim inside the cavity.

When this juvenile leaves, the victim is left alone inside the cavity, surrounded by R1 macaques sitting on the buttress roots around her. The victim sits down and lip-smacks nervously (**Figure S10.4**). SN looks towards her. A juvenile goes inside the cavity and bites the victim on the throat (**Figure S10.5**). While the juvenile was trying to pull from the victim's back, she finally jumps out of the cavity and runs away from R1 (**Figure S10.6**). A couple of juveniles chases her. We searched for the victim, but we could not find her again. Observers did not see any wounded juvenile in R2, and we are uncertain of whether she came back to her group.

#### 29th April 2016: Unknown female and R1

**Available information about the attack:** Four videos of a total duration of approximately 50 minutes. All videos were recorded by Juliette M. Berthier. Below, there is a summary of the event and a full description of what can be seen in the videos.

*Summary*
**Observer/s:** Juliette M. Berthier (JB), Laura Martínez Íñigo (LI), Maura Tyrrell (MT)

**Victim:** Adult female from unhabituated group

**Aggressor group:** R1

R1 was having an encounter with a non-habituated group. R1 surrounded an adult female from the unhabituated group. The rest of her group retreated JB started the recording of events at 10:27.

The victim, an adult female, was curled up between the buttress roots of a tree at the beginning of the observation. Some juveniles and two sub-adult males, Kevin (KK) and Mogwli (MB), looked at the victim. Soon, adult and sub-adult females joined them. Most females that passed by did not pay much attention to the victim. Some examined her. The only exceptions were Vodka (VS), Nuria (NS) and Intan (IU). VS and NS approached the victim for a few seconds, bit her and leave again. IU groomed the victim briefly. KK attacked the female in a couple of occasions at the beginning of the observation. The bulk of the observed attacks came mostly from two unidentified sub-adult females who bit the victim repeatedly, to the extent of dragging her body using their mouths. Several juveniles participated too, examining, biting and even leaning on the female. The victim did not struggle at any point and played dead most of the time. The only exception was when a strong noise was heard, and the victim attempted to stand up. However, as soon as she moved, IU and a juvenile approached the victim, who became still again.

Interestingly, around the middle of the observation, Lucifer (LK), a sub-adult male, approached the victim and sat nearby. While he was there, the attacks ceased almost entirely and the macaques nearby no longer paid attention to her. During this period, LK self-groomed most of the time while keeping body contact with the outgroup female by putting his leg over her head. Later, Martabak (MM), an adult male, approached LK and the victim.

MM smelled the victim and joined LK in that sort of "guarding". Shortly after, MM ran towards a conflict.

Late, Niko (NK) displaced LK and occupied his position. NK did not aggress the outgroup female. Most of the monkeys around did not try to interact with the victim at this point. If any macaque tried, NK threatened them. NK lip-smacked at the victim in a couple of occasions.

Finally, the victim stood up and started walking away slowly before fleeing at full speed. NK, LK and a juvenile chased her. We (LI, MT, JB) tried to find her again, but we were unable. We did not see any injuries on the female from the unhabituated group during the attack.

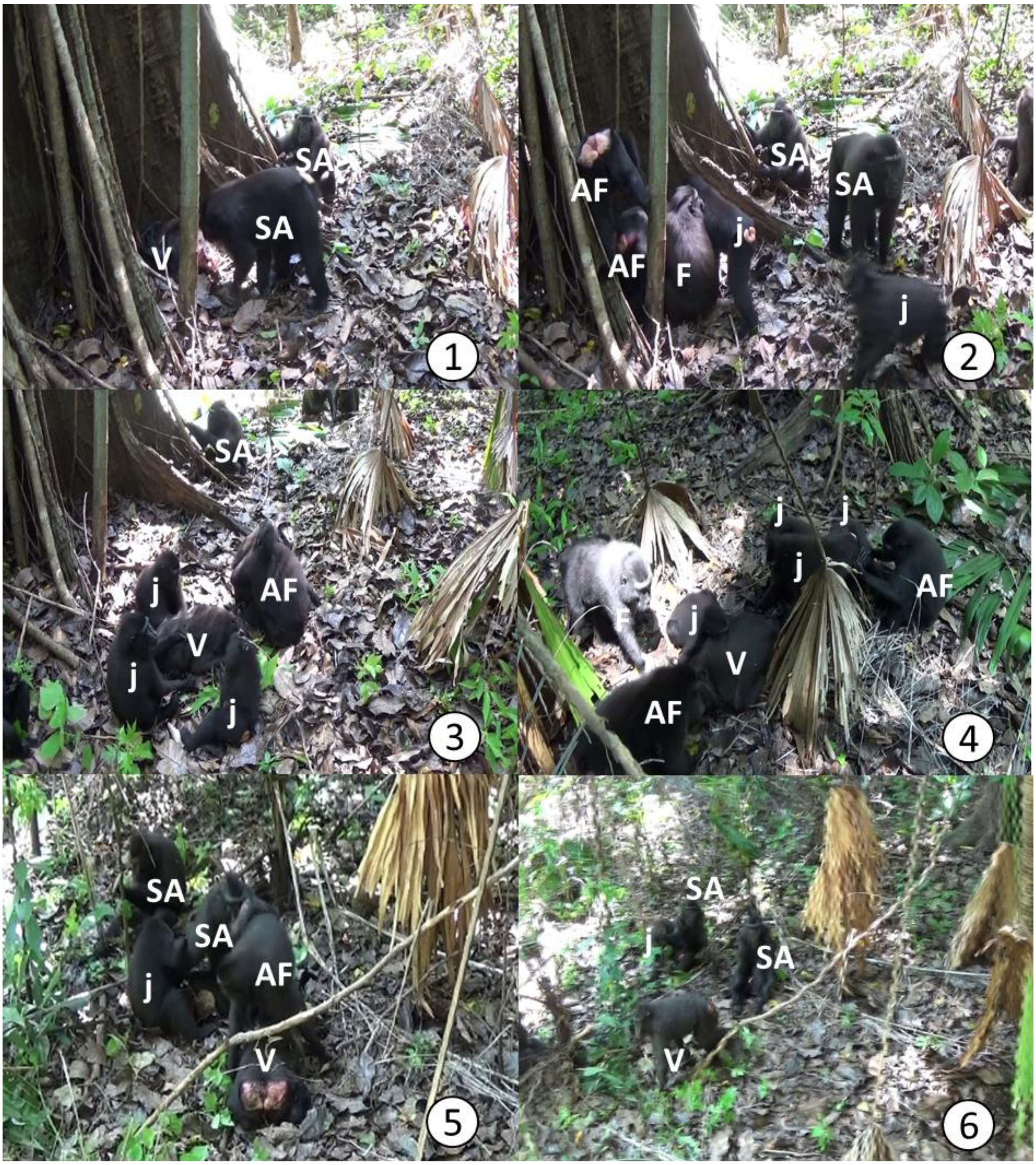

**Figure S11. Members of R1 attacking a female from an unhabituated group the 29th April 2016.** The pictures are numbered chronologically. Age sex classes are labelled as**:** AF= Sub-adult female; F= Adult Female; j=Juvenile; SA=Sub-adult male, V=Victim. Pictures by Juliette M. Berthier

*Full description*

R1 was having an encounter with a non-habituated group. R1 surrounded an adult female from the unhabituated group. The rest of her group retreated JB started the recording of events at 10:27.

At the start, the outgroup female lies motionless between the buttress roots of a tree. Mowgli and Lucifer, both sub-adult males, are in front. Lucifer threats someone outside the video frame and leaves, presumably to chase it. Kevin, another sub-adult male, approaches the victim and manipulates her body, turning it around and examining it. The victim is still. Iris, a sub-adult female, passes by and stops to have a look at the victim. Kevin bites the victim on the flank. Mowgli and Iris leave. Xiro, a sub-adult male, approaches while the victim moves the head to look around. Her breath seems quick. Xiro smells her genitals and then touches her (**Figure S11.1**). The victim curls up, and Kunti, an adult female, approaches her. Kunti touches her and Xiro pulls from the fur of the victim's back several times. A juvenile, Juni, who is an adult female, and a sub-adult female approach the victim (**Figure S11.2**) and Xiro leaves. Then, another juvenile comes close. The macaques cover the view, but it seems that they are all close to the victim, perhaps touching and biting her.

Next, the last juvenile to join the mob, a male, drags the victim toward him, makes her sit and bites her throat. A sub-adult female who we will refer here as AF1, pulls the victim away from the juvenile and bites her. Kunti has left, and Helicopter, an adult female, comes to watch. AF1 seems to bite the victim a couple of times. Ani, an adult female, passes by, looks and continues her way. AF1 leaves and Xiro approaches. He touches the victim. A second sub-adult female, to whom we will refer here as AF2, does the same as Xiro. AF1 comes back and grooms the victim for a couple of seconds before manipulating her body while the AF2 bites the victim's face. AF1 and two juveniles bite the victim too. Caca, a sub-adult female, appears and watches. AF2 bites the back of the victim fiercely. The victim does not struggle at any point but sometimes seems to try to curl up more. Helicopter, who appears again, watches. Caca, AF2 and Helicopter leave. The victim lies motionless facing the ground.

Juni and a juvenile male approach the victim. Juni examines the victim while the juvenile bites her. AF2 displaces the biting juvenile and bites the victim, dragging and shaking her. Xiro approaches and touches the victim. Xiro leaves and a juvenile grooms the outgroup female. Next, AF2 bites her again.

Nihil, an adult female, approaches, smells the female and touches her, almost grooming. She leaves when the af2 comes again and bites the female. Three juveniles examine the female and touch her. Ani approaches, grabs the crest of the victim and pulls a bit from it, apparently to see her face. AF2 tries to affiliate with Ani, who leaves. The three juveniles keep examining the victim. One of them grooms her. Vodka, an adult female, approaches and smells the victim. Juni displaces Vodka and smells the victim.

AF2 comes back, takes a tumble over the victim, and embraces Juni (**Figure S11.3**). Meanwhile, the juveniles keep examining the victim's body. One of them leans on her and grooms her briefly. Ani approaches with Polina, who is another adult female. Polina examines the victim while Ani embraces a juvenile. Leoni, an adult female, approaches the victim, examines her, and smells her. Polina leaves, and Leoni follows her. Nihil approaches the victim and smells her. Nuria does the same in addition to biting the victim. Next, Nuria leaves followed by Nihil. Martabak, an adult male, appears and leaves.

AF1 takes one of the legs of the victim and drags her. Then AF1 pauses before dragging the victim by one of her arms, which AF2 bites. Iris approaches and turns the victim around by pulling from her arms. AF2 bites the victim while some juveniles approach and examine her. Nihil approaches, and then she takes the victim's arm and smells her flank. Two juveniles groom the victim briefly. Next, a juvenile bites the victim's head, and so does AF2 on the face. A juvenile bites the victim´s back. After a pause, AF2 bites the victim's back again. Leoni approaches and smells while a juvenile bites the victim. Leoni touches the victim, whom Nihil smells before leaving. Eight juveniles and Leoni surround the victim. A juvenile grooms the victim's back. Leoni leaves. Iris approaches the victim, smells her, and drags her. A juvenile bites the female's back. Iris retreats and Kunti approaches. Uhnil, a sub-adult female, comes near the victim and smells her. AF2 approaches and bites the female several times. Then the AF1 approaches and does the same. AF2 bites the victim's leg and drags her. AF2 leaves and a juvenile grooms the victim. AF2 comes back and bites the victim. AF2 leaves, comes back again, and bites the victim before dragging her. Then, AF2 pauses, bites the victim once more and leaves. Five juveniles and AF1 are still around the victim at this point.

Intan approaches and smells the victim. A juvenile touches the victim and lies by her side. Another juvenile bites the victim playfully. AF2 approaches and bites the victim's mouth. Then, AF2 bites the victim's back and drags her. AF2 bites the victim once more, and a juvenile does the same (**Figure S11.4**). AF2 bites the victim's face again and drags her. Then Vodka bites the upper part of the victim's head and drags her. A juvenile bites the victim. After a pause, the juvenile and AF2 bite the victim again. Next, AF2 displaces the juvenile to bite where it was biting and takes a tumble over the victim.  Two juveniles groom the victim. One of them leaves, and the other continues. Kunti approaches and smells the female. A long time passes with Vodka, Kunti, AF1, AF2 and several juveniles around the female without attacking her. Then AF2 and a juvenile bite her again.

Lucifer approaches, sits and briefly grooms the victim. Kunti leaves. Lucifer stays by the victim grooming himself. AF2 approaches but retreats when Lucifer looks at her. Later, a juvenile tries to approach but moves away when Lucifer threatens her. After, Lucifer lunges to a juvenile nearby, who goes further. When he does that, the victim reacts, straitens a bit and looks around. Immediately afterwards, a juvenile and Intan approach the victim and examine her. Then AF2 bites the victim on the face and drags her. A juvenile grooms the victim and bites her. Intan smells the victim's anal area. Later, a juvenile put itself over the victim, tries to turn her around and bites her. Next, a juvenile grooms the victim briefly. Only Lucifer, Intan and two juveniles are within 1 meter of the victim. Lucifer approaches the victim displacing the juveniles that were close to her. Lucifer sits by the victim, with one of his legs over her head. Both juveniles and Intan leave and Lucifer grooms himself. Martabak approaches and smells the victim before sitting close to her. Lucifer pets the victim's back.

AF1 approaches and examines the victim. She sits for some seconds and then leaves the proximity of the female. For a long time, only Lucifer and Martaback stay less than a meter from the victim. Lucifer grooms himself and Martaback looks around. The victim lies playing dead. After a while, Martabak runs towards a conflict out of sight. AF2 approaches and looks at the victim before sitting within a meter. A bit later, a juvenile male joins AF2. Lucifer still has his leg over the victim's head. Then Niko approaches and smells the victim, displacing Lucifer. Lucifer sits within 2 meters of her and Niko. The AF2 smells the victim, bites her quickly in the leg and retreats. Next, AF2 gets closer again to the victim and smells her anal area. Niko smells and touches the female. Sometime later, AF2 leaves the proximity of the female.

A juvenile approaches, presents to Niko. Niko presents to the juvenile for grooming and the juvenile grooms him. The juvenile male who arrived with AF2 smells the hindquarters of the victim. Another juvenile male approaches and does the same. AF2 does as well. Next, Niko threats them and the juveniles retreat. AF2 touches and grooms the victim, ignoring Niko's threat. Then she leaves the close proximity (<1m) of the victim.

A couple of minutes later, Caca, a sub-adult female approaches. She presents to Niko while she stands over the female (**Figure S11.5**). Niko examines Caca but rejects her. Then Caca grooms the victim, briefly before leaving.

Niko lip-smacks to the victim. A juvenile approaches. Niko lies down and puts his face close to the victim. The juvenile who was grooming Niko stops and leaves. Niko lip-smacks and the other juvenile leaves.

Niko continues lip-smacking at intervals to the victim. Finally, the female stands up and slowly starts to move away with silent bared-teeth, before fleeing (**Figure S11.6**). Niko, Lucifer and a juvenile run behind her. We could not find her again. We did not see any injury on her during the attack.

#### 18th July 2016: R1 attacks five individuals from PB1

At 13:09, we (Laura Martínez Íñigo (LI) and Juliette M. Berthier (JB)) perceived that many individuals of R1, our focal group that day, ran towards the beach, to the south. I (LI) followed them and saw that PB1 was there, fleeing towards the east. R1 run perpendicular to them. Most of PB1 managed to escape. However, several individuals were kept by small groups of R1 macaques. JB and I followed as many cases of such harassments as possible as reported below. We recorded 5 but there was at least one more that I observed briefly while recording another attacked.

**Available information about the attacks:** Four videos, two by Laura Martínez Íñigo and two by Juliette M. Berthier. We recorded simultaneously different attacks, generating a total of an hour of footage recorded within 40 minutes. Parts of these videos can be watched in **Video S3**. The description of each event is summarized and at fully transcribed below.

*Emping and her infant*

Summary

**Observer/s:** Laura Martínez Íñigo (LI) and Juliette M. Berthier (JB)

**Victim:** Adult female (Emping) and her infant, from PB1

This one was the first mob that I (LI) approached was a pool filled with water on the rocks by the beach. Surrounding the pool, I found the adult male Martaback (MM), the adult females Tuti (TS), Ola (OU), Polina (PS), Yane (YS), Vodka (VS) and Leoni (LS) as well as two juveniles. They were all from R1. Inside the pool, there was an adult female from PB1, Emping, with her infant.

For about 10 minutes, several juveniles, LS and OU grabbed Emping, pulled from her arms and bit her. Some juveniles tried to touch the infant as well. Emping did not resist nor move except to cover her infant once it was exposed. Emping defecated several times inside the pool during the attack. Her infant swam in the puddle but remained close to her mother and grabbed her fur when possible. Eventually, with the pool still surrounded by several R1 monkeys, Emping jumped out from the pool and fleed carrying her infant.

However, the subadult female Caca (CB) and several juveniles chased Emping and her infant. They caught her and formed a huddle around her, apparently aggressing both macaques. Soon, Emping jumped into the water and swam to a nearby rock while her infant stayed behind. A subadult female and several juveniles attacked Emping, and she ended up swimming to another rock. Meanwhile, her infant was bitten and licked by the subadult male Lucifer (LK). The infant protested every time anyone touched her and moved towards the forest whenever she was left alone. She lip-smacked whenever another monkey looked at her. Nuria (NS, adult female) suddenly approached and bit the infant violently. Then a wave broke, and everyone around the infant left, leasing her.

While the infant kept moving slowly towards the forest, her mother was attacked. Ani (AS, adult female), Kevin (KK, subadult male) and CB chased Emping, who jumped onto the sea and swam away.  I continued following another coalitionary attack on a juvenile female (see below).

Emping reached the place where JB was recording the aggression on Gluten (GA, subadult female from PB1, see below). There, Mowgli (MB, subadult male), LK and a juvenile were aggressing GA. Interestingly, when MB and the juvenile went out of site and LK attacked Emping, she resisted. Emping released herself by struggling and screamed while lunging at LK. Then she retreated and went behind GA. LK grabbed Emping again, and she resisted, screaming and even hitting LK.

While they are fighting, MB approaches, grabs Emping's crests and pulls, making her fall back. Emping's behaviour on this occasion is strikingly different from that in the pool, where she did not defend herself.

Soon after, the three macaques from R1 macaques left, and Emping swam away. Once Emping reached the nearby rocky beach, several juveniles chased her, and she fled towards PB1 and rejoined them. Nobody reported injuries on Emping.

Emping's infant reached the forest. There, she Adinda (AU), an adult female from R1, protected the infant and carried her. However, the infant died one day later. AU carried

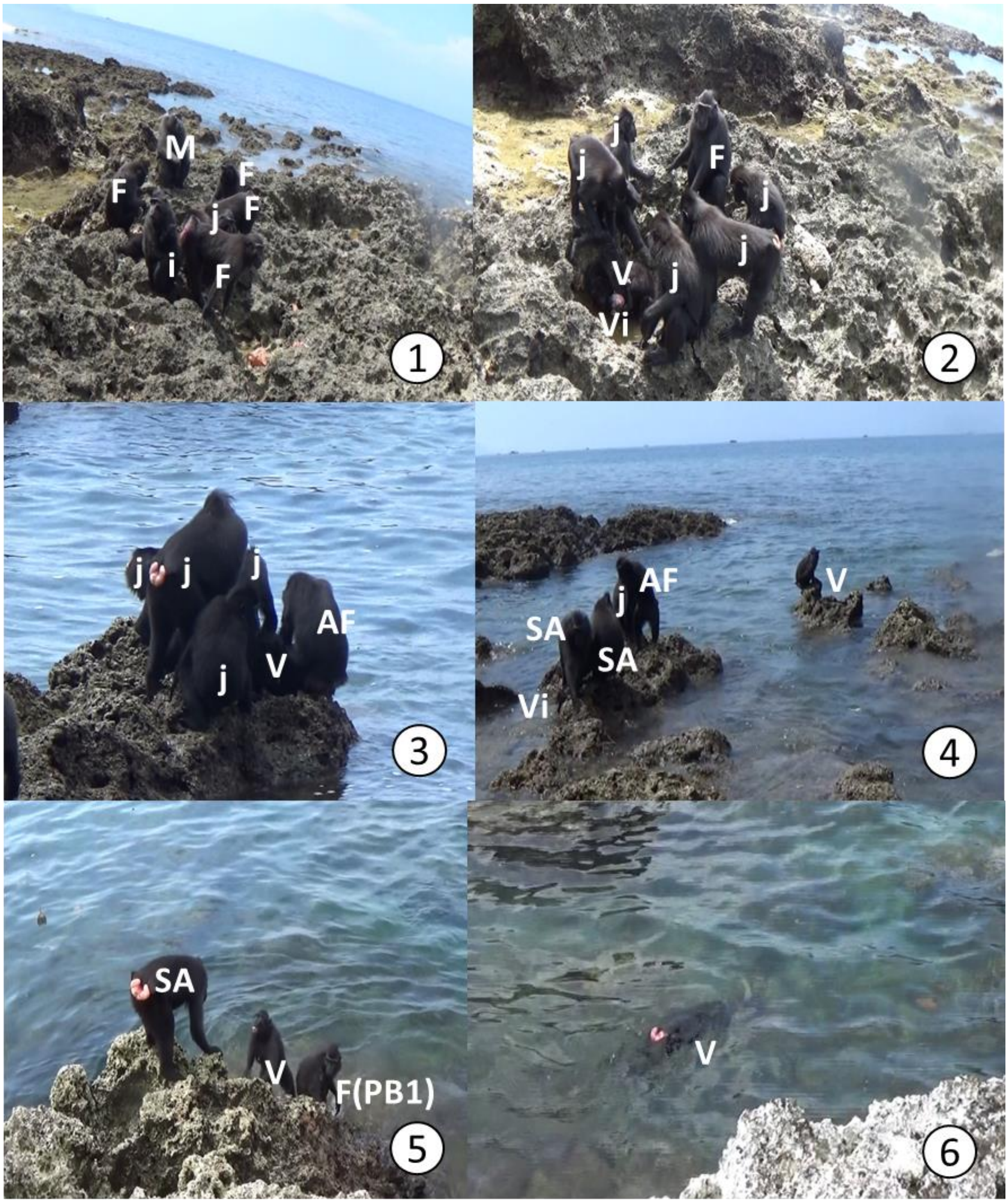

**Figure S12. Members of R1 attacking an adult female from PB1 (Emping. EA) and her infant the 18th July 2016.** The pictures are numbered chronologically. Age-sex classes are labelled as: AF= Sub-adult female; F= Adult Female; j=Juveniles; SA=Sub-adult male; V=Victim ; Vi= Victim’s infant ; M=Adult Male. Pictures by Laura Martínez (1-4) and Juliette M. Berthier (5-6).

Full description

This one was the first mob that I (LI) approached was a pool filled with water on the rocks by the beach. Surrounding the pool, I found the adult male Martaback (MM), the adult females Tuti (TS), Ola (OU), Polina (PS), Yane (YS), Vodka (VS) and Leoni (LS) as well as two juveniles. They were all from R1. (**Figure S13.1**) Inside the pool, there was an adult female from PB1, Emping, with her infant.

Shortly after I arrive, Yane, Vodka and others run towards another individual from PB1 (Maybe Gluten? see below for details on the attack against her). Leoni is then the only macaque from R1 staying by the pool. Leoni bends to reach Emping and bites her. Ola comes back with her infant and a juvenile. Ola bites Emping as well. Kunti, Martabak and a juvenile approach too. The observation is then briefly interrupted when I detect another coalitionary attack, and I call JB to cover it.

The attacks on Emping and her infant stop for some time. Emping seems paralyzed, crouching inside the pool. Ola and her infant baby are at the edge of the pool. Some juveniles are visible, and they observe what is happening on the other side of the beach. Emping's infant climbs to her mother's back, and then Leoni and two juveniles approach. Ola leaves with her infant. A juvenile and Leoni reach out to Emping and the infant and touch them. The other juveniles approach and watch the infant. One of these juveniles bends and lip-smacks at the infant. Emping defecates several times. Other than that, she does not struggle when the macaques of R1 grab her. Leoni touches her gently and then she grabs Emping's hand and pulls. This action forces Emping to sit down on the pool. Emping looks around. A juvenile grabs the fur of Emping's back and pulls towards itself. Next, the juvenile pulls from Emping's arm and bites it. Emping does not resist. Another juvenile touches Emping's head. About 1-2m from the pool, Vodka embraces Ola, and then Leoni approaches them and embraces Vodka.

Meanwhile, a juvenile bites Emping. When the embrace is over, Vodka approaches Emping vocalizing and pulls the fur from Emping's back, forcing Emping to look at her. Leoni leaves. A juvenile pulls from Emping's arm several times. Emping does not resist but breathes quickly. A juvenile pulls Emping's arm again while another pulls the fur of her back until they turn her around, exposing the infant (**Figure S13.2**). Next, one of the juveniles rubs the infant's face and pulls from Emping's and her infant's arms. When the juveniles pulling, Emping curls, hiding the infant under herself as she was doing before. Both of these juveniles were females. One of them pulls from Emping again. Leoni pulls too, and a juvenile seems to bite Emping. Around 13:17, the observation stops for a moment to watch another coalitionary aggression happening far away, where Mowgli (sub-adult male) and a sub-adult female seem to be aggressing a juvenile. I continue observing the case of Emping and her infant.

Two juveniles and Leoni grab Emping's fur and pull from her to different sides, vocalizing. When they stop, Emping jumps out of the pool and runs away towards the sea, chased by several juveniles and carrying her infant. Caca catches Emping and bites her. Emping shows no resistance. Several juveniles approach and seem to touch Emping and her infant. One of the older juveniles seems to be biting either Emping or her infant (**Figure S12.3**). Another sub-adult female joins the mob. Caca bites Emping or her infant, and some juveniles seem to be doing the same. The infant screams. Then Emping jumps to the water and swims to the next rock, leaving her screaming infant behind. While several individuals surround Emping, Lucifer takes the infant and pushes it onto the rocks while the infant protests.  Emping is left alone (**Figure S12.4**). Lucifer bites the infant's back, which makes her complain complains. A sub-adult female seems to lip-smack towards Emping. Then an old juvenile jumps towards Emping, grabs her crest, and pulls her into the water.

Meanwhile, Lucifer licks Emping's infant. Then, once Emping gets out of the water, an adolescent sub-adult female jumps to the rock Emping is on, grabs her crest, and pulls down. Emping does not resist.  Lucifer keeps licking the infant's back while grabbing it with both hands. The infant lip-smacks at Mowgli, who seems to lip-smack back. Lucifer bites the infant's back again with the side of his mouth. A juvenile approaches the infant and seems to bite it as well.  Both Lucifer and the juvenile release Emping's infant, who jumps to another rock towards the forest. Lucifer follows the infant and presses it against the ground. Next, Lucifer bites the infant's back superficially, and the infant complains. Meanwhile, Vodka and a juvenile are biting Emping, who jumps to the water and swims to another rock.

Lucifer bites the infant superficially and then holds it back by grabbing its leg. Vodka approaches the infant and lips-smacks at it. Then Lucifer bites the infant while it complains, and Vodka vocalizes (**Figure S13.1i**). Lucifer pulls the infant off the ground; Vodka touches it. Nuria approaches and bites the infant on the neck violently (**Figure S13.2i**). A wave breaks and makes all the monkeys on sight to run away towards the forest.

The infant lags and is caught by Lucifer, who bites it again. Then Lucifer leaves, and the infant seems dead (**Figure S13.3i**)., but soon it stands up and walks towards the forest (**Figure S13.4i**).

Meanwhile, its mother is coming out of the water and alone, looking towards the forest. The infant keeps moving slowly towards the forest and lost-calling, sitting, and lip-smacking if a monkey of R1 looks at her. Then Caca, Kevin, and Ani run towards Emping, who jumps into the sea before they can catch her and swims away. Shortly after, I detected an attack on a juvenile from PB1 (see below) and started recording it.

Eventually, Emping arrives where JB is recording the attack on Gluten (see below) around 13:25. Emping swims up to the rocks where Gluten is and sits close to her. The only members of R1 near them are Lucifer, Mowgli, and a juvenile. Soon after reaching the rocks, Lucifer and the juvenile threat Emping, and she does silent bared-teeth. The juvenile approaches her and seems to aggress her. He repeats the operation, hitting Emping's head with his hand flat. The juvenile leaves, and so does Mowgli. Then Lucifer approaches Emping. He grabs her by the crest and pulls down to bite her neck. She screams and struggles and manages to release herself (**Figure S12.5**) and go behind Emping. The juvenile and Mowgli appear again, and Lucifer grabs Emping again and bites her several times while she screams and struggles. Then Mowgli approaches and hits her. Lucifer releases her, and Mowgli grabs her crest and makes her fall while she screams. After that, both sub-adults retreat, and so does the juvenile after lip-smacking at Emping. The sub-adults leave, and shortly after, Emping jumps into the water and swims and dives away. Once she arrives on the nearby rocky beach, she is chased by juveniles and runs towards PB1. She reached them and joined them successfully. No injuries were reported for her.

Emping´s infant reached the forest where Adinda, an adult female from R1, started carrying her. Adinda protected the infant if someone tried to approach her. However, the infant died one day later, without conspicuous signs of violence. Adinda carried the body one more day after the dead of the infant. We recovered the body. However, it had been dragged after the dead of the infant and was not possible for us to assess the *pre-mortem* wounds.

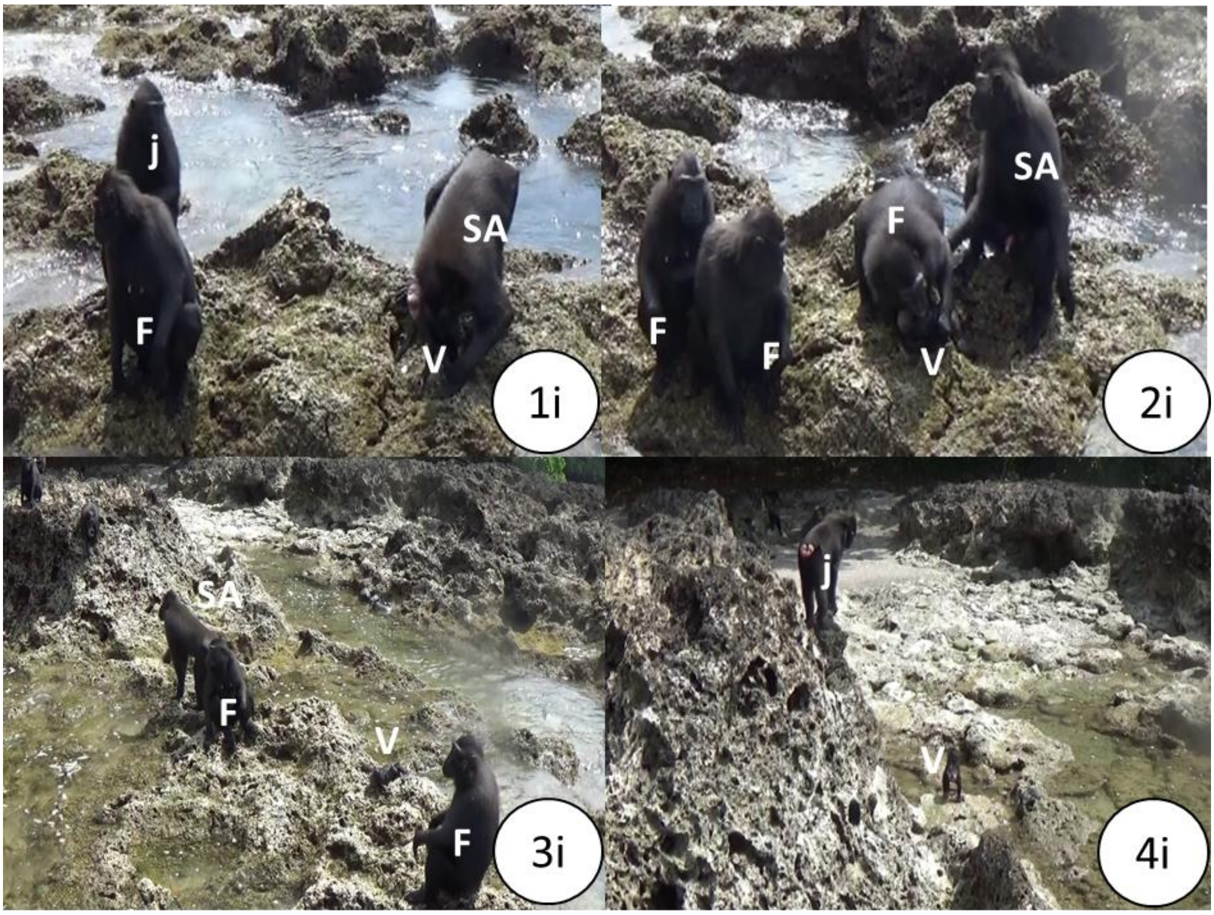

**Figure S13. Members of R1 attacking an infant from PB1 after separating her from her mother (Emping, EA), the 18th July 2016.** The pictures are numbered in chronological order F= Adult Female; j=Juveniles; SA=Sub-Adult male; V=Victim. Pictures by Laura Martínez Íñigo.

*Gluten*

Summary

**Observer/s:** Juliette Morgan Berthier (JB)

**Victim:** Sub-adult female (Gluten) from PB1

JB arrived at a mob where Adinda (AU, adult female), Helena (HS, adult female), Caca (CB, sub-adult female), Kevin (KK, sub-adult male), Proboscis (OM, adult male), and several juveniles are surrounding Gluten (GA, sub-adult female from PB1). OM bent over GA and left shortly after. GA soon managed to escape, but KK and two juveniles chased her. They aggressed and touched GA. At some point, Solo (SK, adult male) approached GA. SK seemed to direct some facial expressions at GA (not visible). She grimaced. When SK attempted to get closer, GA screamed and avoided him. Then SK retreated and left. KK came back and aggressed GA several times. GA resisted the attack and hit him. A juvenile aggressed GA well. Mogwli (MB, sub-adult male) appeared with Polina (PS, adult female), who bit GA.

Shortly after, Emping (see above) appears swimming. Lucifer (LK, sub-adult male), MB, and a juvenile are now the only R1 macaques near them. The juvenile grabbed GA and bit her. Later the three macaques from R1 retreated. Emping swam away, and GA fled towards PB1 and out of sight. She joined PB1 successfully, and no injuries were reported for her.

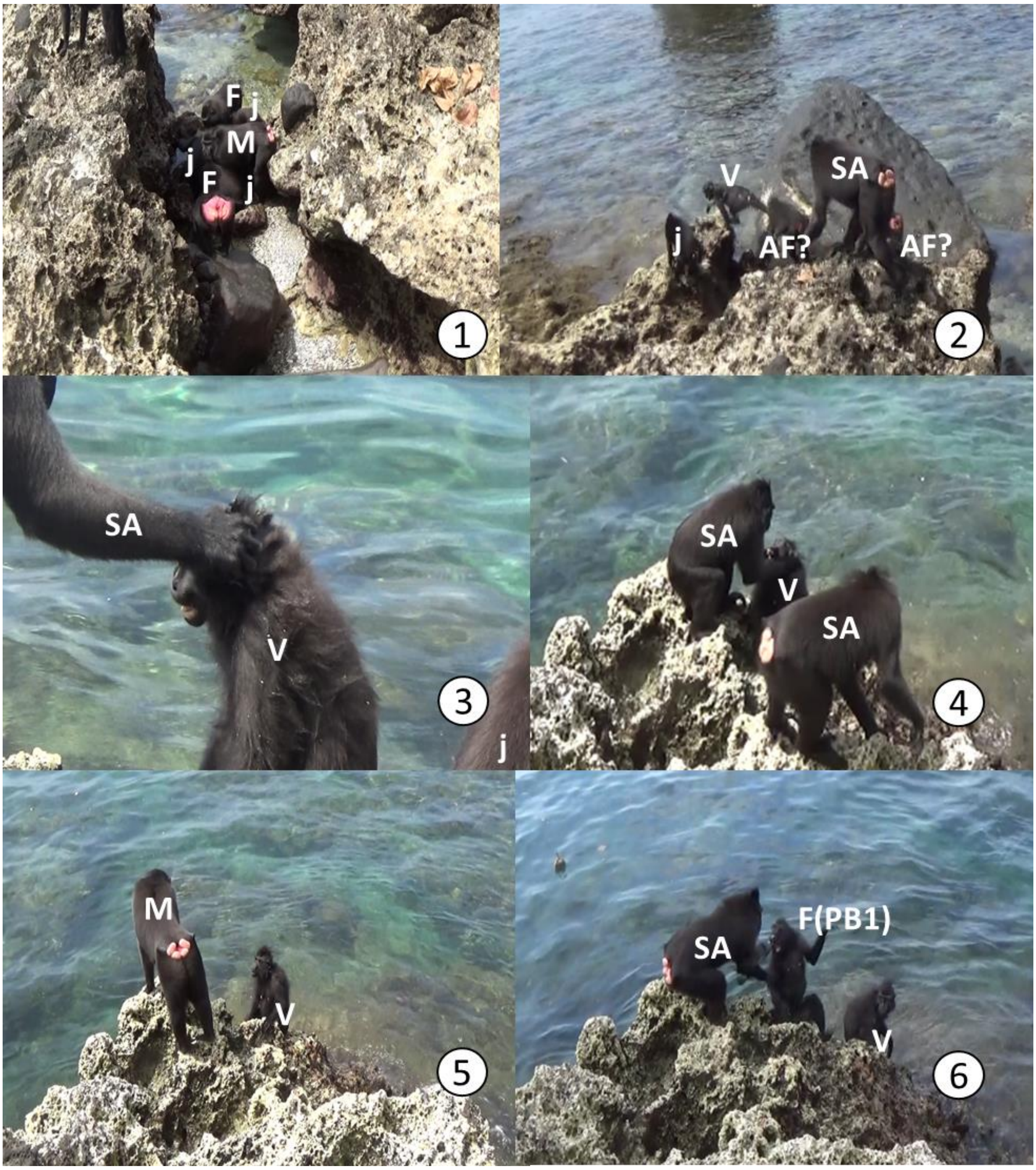

**Figure S14. Members of R1 attacking a sub-adult female from PB1 (Gluten, GA) the 18th July 2016.** Pictures are numbered in chronological order. Age-sex classes are labelled as AF=Sub-adult female F= Adult Female; j=Juveniles; SA=Sub-Adult; V=Victim; M=Male. F(PB1): Adult female from PB1 (Emping, EA). Pictures taken by Juliette M. Berthier

Full description

At 13:11, JB arrives at a mob where Adinda, Helena, Caca, Kevin, Proboscis, and several juveniles surrounded Gluten (**Figure S14.1**). It seems that at least the two juveniles are attacking her, either pulling her hair or biting (they are on top of her and is not visible). Proboscis is bent towards Gluten as all others. Proboscis leaves, and the others diminish their attack. Gluten escapes (**Figure S14.2**). Kevin and two juveniles chase Gluten towards the sea and surround her. She half-open mouths, and Kevin touches her head several times (**Figure S14.3**). Then Kevin walks away, and a juvenile approaches Gluten and touches her. Kevin grabs Gluten and pushes her head down. She screams and struggles. Caca approaches and bites Gluten. Then Caca leaves, and Kevin approaches again, grabs Gluten, and forces her to look at him (Figure S14.4), after which he retreats. Kevin approaches again, touches Gluten head, and leaves.

Solo approaches, Gluten opens-mouths at him and avoids him. Solo stands close to Gluten and seems to direct facial expressions at her (but is with his hindquarters towards the camera, it is impossible to see his face) (**Figure S14.5**). Solo lunges at Gluten, and she screams and avoids him. Then Solo leaves, Kevin approaches, stands, yawns, and leaves again. Shortly after, Kevin repeats the same. Then he comes back, grabs Gluten, and tries to bite her while Gluten screams and struggles. There is no one else nearby. Kevin leaves and approaches Gluten again, grabs her by the crest and pulls towards her back. Then Kevin retreats again. A juvenile approaches Gluten, examines her, and when he tries to bite her, she screams, struggles, and hits it. Then the juvenile retreats, and Kevin pushes her head down violently. Then he grabs Gluten's face and makes her look at him. Then he bites her while she screams and struggles.

After a moment covering the attack on Yams (see below), the observation continues with Gluten, who screams as Mowgli approaches, threatening her. Gluten retreats. Polina and Mowgli approach. Polina grabs Gluten's face and bites her. The juvenile bites Gluten too. Then Polina leaves. After a while, a juvenile male goes close to Gluten, turns her around, and bites her back softly. Then he retreats. Lucifer approaches her and directs some facial expressions at her, but it is impossible to see which ones.

At this point, Emping arrives swimming and sits close to Gluten (see above). While Lucifer attacks Emping, Gluten retreats slightly and looks towards them (**Figure S14.6**). A juvenile male approaches Gluten, grabs her, and bites her while Gluten struggles. Mowgli and a juvenile retreat, leaving Emping and Gluten alone with a male juvenile. The juvenile retreats shortly after. Emping escapes swimming, and then Gluten flees towards PB1 and out of sight. She joined them successfully, and no injuries were reported for her.

*Female juvenile from PB1*
Summary

**Observer:** Laura Martínez Íñigo (LI)

**Victim:** Juvenile female from PB1

After Emping was separated from her infant, who was going towards the forest, I (LI) encountered another mob attacking a juvenile female from PB1 (13:23). The juvenile from PB1 screamed while a juvenile from R1 and Nuria (NS, adult female) bit her. Two adult females (Intan, IU, and Juni, JU) were bystanders, together with some more juveniles. This means that they were within 2 meters, but they did not interact with the victim. The juvenile from PB1 struggled and managed to free herself and run, but was soon trapped again. She screamed and did silent bared-teeth to the macaques of R1, who tried to approach her. Caca (CB, sub-adult female) and Kevin (KK, sub-adult male) were the most aggressive. They followed the juvenile from PB1 from rock to rock in which she tried to escape and swim away from them. Whenever they reached her, they bit her, hit her, and pushed her into the water.  Ani (AS, adult female) bit her once as well. The juvenile from PB1 ended up swimming up to the rocks in which Emping (EA, adult female from PB1) and Gluten (Sub-adult female from PB1) had been before. There, the juvenile from PB1 was attacked there by several juveniles, who hit her and pushed her into the water. In the end, she managed to re-join PB1. Four juveniles in PB1 were reported to have injuries that day, but since we could not tell them apart, we do not know whether this specific juvenile was wounded.

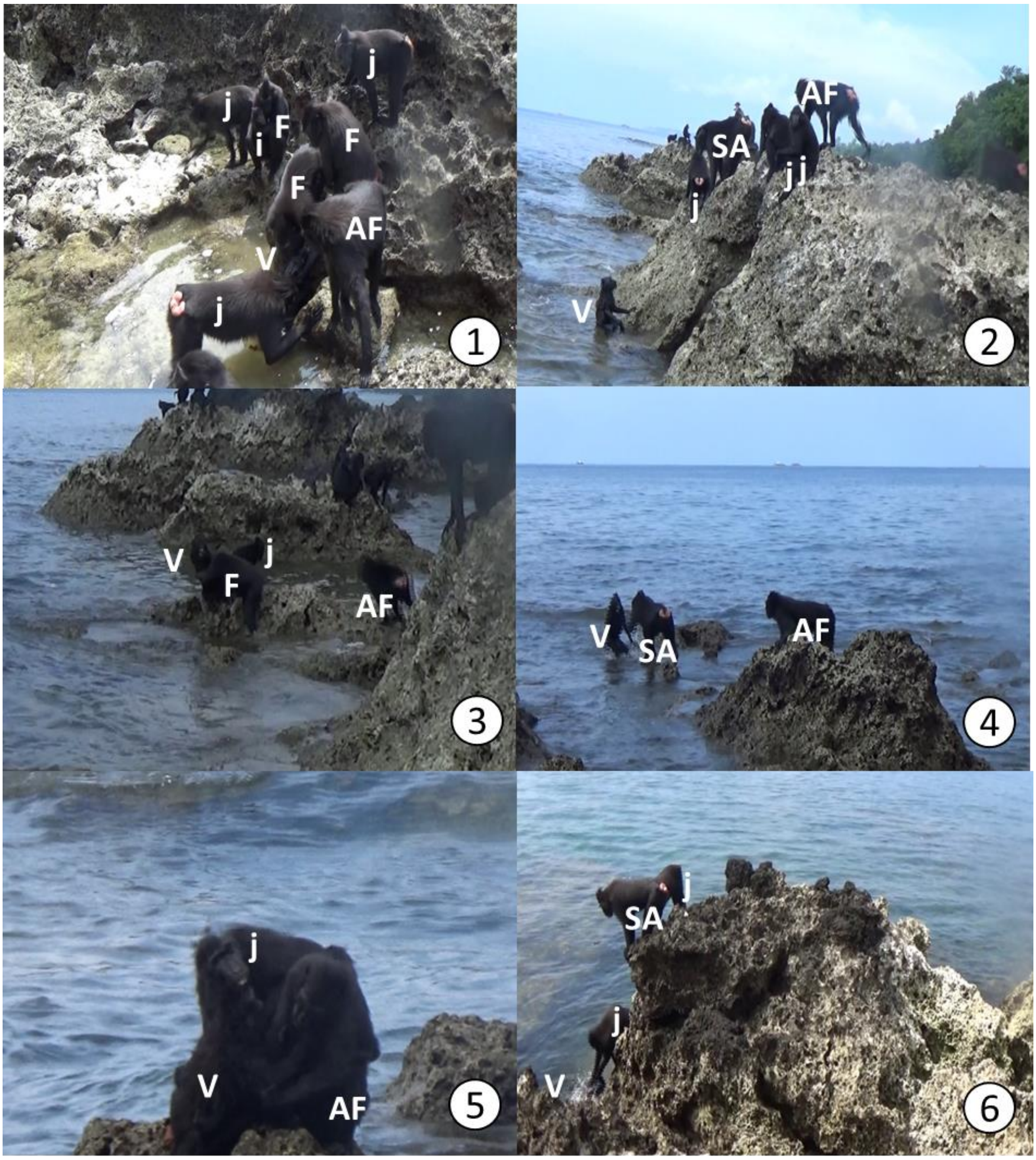

**Figure S15 Members of R1 attacking a juvenile female from PB1 the 18th July.** Pictures are numbered in chronological order. Age-sex classes are labelled as: AF=Subadult female F= Adult Female; j=Juveniles; SA=Sub-Adult; V=Victim ;i=Infant. Pictures by Laura Martínez Íñigo

Full description

After Emping was separated from her infant, who was going towards the forest, I (LI) encountered another mob attacking a juvenile female from PB1 (13:23). The juvenile screamed while a juvenile and Nuria (NS, adult female) bit it (**Figure S15.1**). Two adult females (Intan, IU, and Juni, JU) were bystanders, together with some more juveniles. This means that they were within 2 meters, but they did not interact with the victim. The juvenile was most likely a female since the groin is visible several times in the video, and no sexual organs are seen (Males have pink genitals, easy to spot since they are born). Two juveniles and Caca (CB, sub-adult female) grab the juvenile from PB1 while trying to escape. The juvenile from PB1 screams and struggles. She manages to release herself and runs away a few meters, but she is chased by several juveniles and CB. Then they pushed her into the water (**Figure S15.2**). There, the juvenile from PB1 has a break from the attacks, since R1 seems to be unwilling to venture in the water. The macaques from R1 stand lip-smacking at her and approach her, trying not to touch the water. At some point, the juvenile from PB1 sits on a rock. Then, a juvenile from R1 comes and grabs one of her arms. Ani (AS, adult female) then approaches and bits the juvenile from PB1 on her arm. The juvenile from PB1 screamed while bitten (**Figure S15.3**). A sub-adult female approaches and tries to grab the juvenile from PB1. However, the juvenile from PB1 escapes by swimming into the water until she reaches another rock on the surface. Then she moves to another rock. There, a juvenile from R1 embraces her. The juvenile from PB1 tries to avoid the embrace. Then the juvenile of PB1 is alone for some moments. Then CB jumps to the rock the juvenile from PB1 is on. Kevin (KK, sub-adult male), follows CB. The juvenile from PB1 moves away from the rock, goes into the water, and screams. KK jumps up to where the juvenile from PB1 is, pushing her towards a deeper place. The victim screams and KK hits her, making her sink (**Figure S15.4**). The juvenile tries to swim toward another rock, but KK hits her again, pushing her into the water. The juvenile from PB1 swims away from CB and KK. CB approaches her, but then the observation is interrupted to look for Emping's infant and the detection of another mob. When the observation on the juvenile from PB1 continues, she is on a rock surrounded by water. CB and an old juvenile, possibly Qubes (QB), jump onto the rock and bite the juvenile from PB1 (**Figure S15.5**). The juvenile from PB1 struggles and jumps to the water. KK leaves, and the juvenile from PB1 swims to the rock CB is on. The juvenile from PB1 sits close to CB and lip-smacks. Then CB leaves, and the juvenile stays on the rock. The observation is then interrupted to follow the one against Yams (see below) and only checked at intervals. During one of those, Lucifer (LK, sub-adult male) and a juvenile direct facial expression towards the juvenile from PB1, but it is difficult to see which ones. The juvenile from PB1 swims towards another rock and screams when several juveniles threaten her at her arrival. A juvenile grabs the victim and hits her (**Figure S15.6**). The victim screams and lip-smacks. Another juvenile approaches, and lip-smacks at the victim. Next, this juvenile from R1grabs the juvenile from PB1 and pulls from her while the victim screams. The observation is then interrupted again to check for Emping's infant. When I come back, I see (and record) blood on the rocks around the juvenile from PB1, which is now alone. A juvenile from R1 comes, grabs her, and the juvenile from PB1 ends up in the water again. It comes up, and the juvenile of R1 leaves. The observation was then interrupted to check others, and shortly after all PB1 retreated. The juvenile was seen fleeing towards them. Four juveniles in PB1 were reported to have injuries that day. However, since we could not tell them apart, we do not know whether this specific juvenile was wounded.

*Yams*

The information available from this event was limited and a summary was deemed unnecessary.

Full description

**Observer:** Laura Martínez Íñigo (LI) and Juliette M. Berthier (JB)

**Victim:** Female from PB1 (Yams, YS)

The first recordings of this attack were made by JB at 13:20, while she was covering the attack on Gluten (GA, sub-adult female from PB1). At that point, Yams (YS, an adult female from PB1) is upon a rock surrounded by water while Kevin (KK, subadult male) and Lucifer (LK, subadult male) threat her from the other side of the water (**Figure S16.1**). Then JB continues with the observation on Gluten. At 13:23, JB records how Gina (GU, adult female), Caca (CB, sub-adult female), and a juvenile surround Yams. Caca bites Yam's arm while Gina pulls from Yam's crest. Yams does not resist (**Figure S16.2**). This is the last observation from JB.

While following the events happening to the juvenile of PB1, around 13:30, I (LI) detected four juveniles, an adolescent female, Ani (AS, adult female), and Mowgli (MB. Sub-adult male) around Yams. She was sitting down and paralyzed. Mowgli grabs Yams' forehead and pulls backwards. She does not resist but displays silent bared-teeth (**Figure S16.3**). Mogwli stops. Next, Ani and a juvenile touch Yams briefly, after which Ani stays holding Yams' hand. Caca approaches the mob while a juvenile observes Yams' face intently. Yams teeth-chatters towards Mowgli, whose face is not visible. Then Ani grabs the fur of Yams' forehead and pulls backwards. Yams does not struggle and curls up when Ani stops pulling. Then the observation is interrupted to check on the juvenile of PB1. When observing Yams again within the same minute, only Ani and Mowgli are still around her. Yams sits between them, apparently paralyzed. Shortly after, Yams seems to recover conscience and looks around, detecting where some members of PB1 are. Then she starts walking towards them with silent-bared teeth. Ani grabs her and forces her to turn her head towards her. Then Ani releases Yams, who jumps and flees (**Figure S16.4**). Some juveniles chase her for a few meters, but then they stop once she reaches an area where some members of PB1 are (Codot, adult male, Mr.I, adult male, Fonz, sub-adult male).

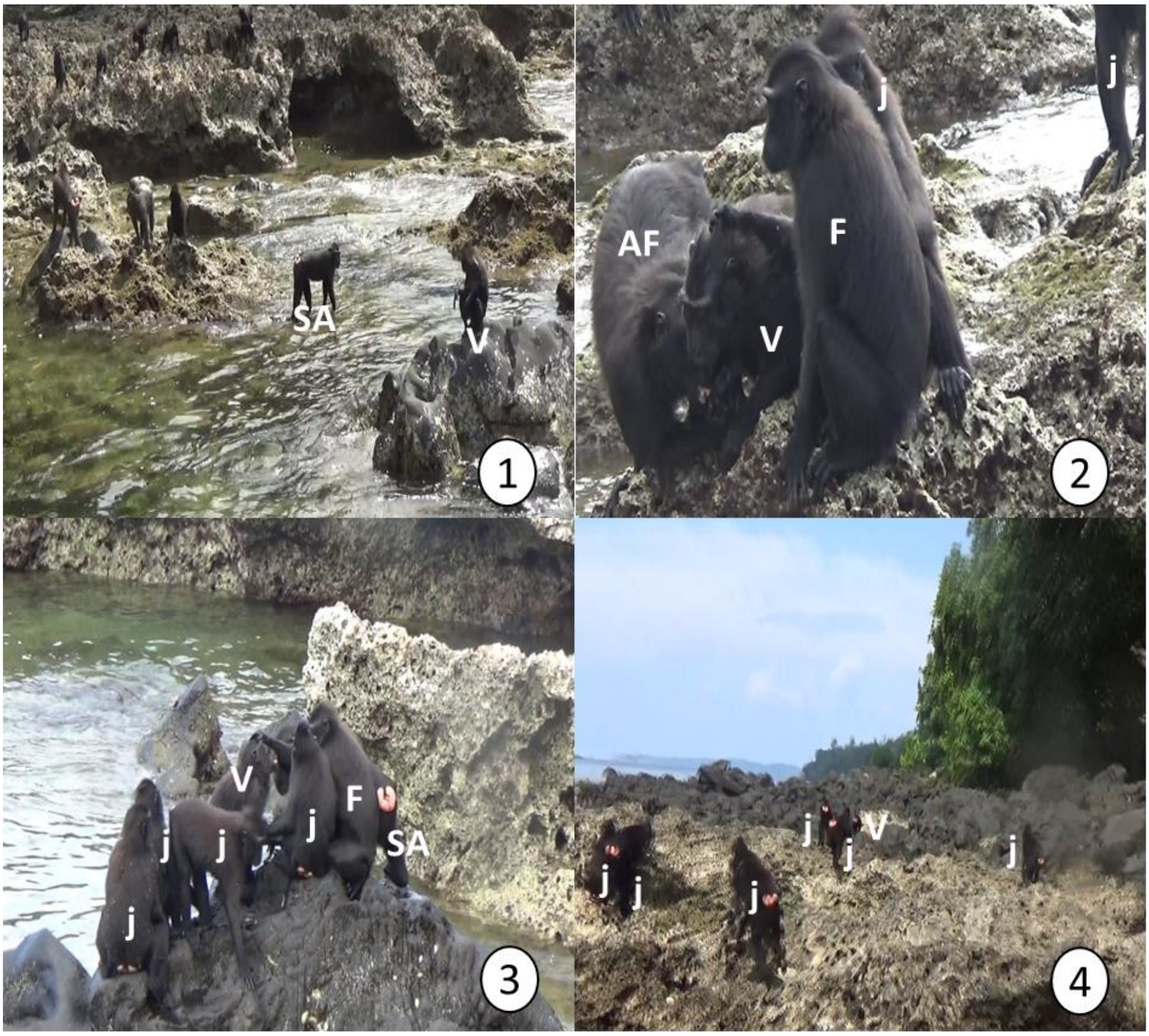

**Figure S16. Members of R1 attacking an adult female from PB1 (Yams, YS) the 18th July 2016.** Pictures are numbered in chronological order. Age-sex classes labelled as: AF=Sub-adult female; F= Adult female; j=Juveniles; SA=Sub-adult male; V=Victim. Pictures by Juliette M. Berthier (1-2) and Laura Martínez Íñigo (3-4)

#### 19th January 2019: R1 attacks adult female with infant from PB1B

**Type of information available:** Field note

**Observer/s:** Unknown

*Field notes*

"*Around 15.12h, group R1 approached group PB1B and thus initiated an inter-group encounter. One of the females of PB1B, Lucia (LA) carried her 7-week old male infant (LA1B) by that time when she was approached by two subadult males from R1. One of the subadult males attacked LA who fought back. During this, the male bit the infant, who was then lost by LA. In the meantime, two additional subadult males had appeared on the scene and the four males chased LA up a tree, where LA stayed until the males had left. Her infant was bleeding from two wounds, one on the left side of the head and one on the left side of the neck, which seemed to stem from two independent bites, and died 2h later.*

*Additional information:*

*On that specific day, LA and another male from her group, Fatman (FM), were found with broken limbs in the early morning. Lucia had her right hand and her left leg broken and was not properly able to walk, which for sure made her an easy target. We believe that they must have sat on the same tree during the night, which then must have broken.*"

#### Discussion on variation in gang attack frequency in crested macaques

Previous researchers working with our study population in 1993-1994 did not observe instances of severe or lethal gang attacks (M. Kinnaird, pers.comm.). However, they saw individuals of one group holding individuals of another back during encounters, which is similar to what we observed during most gang attacks. It is feasible that an increase in population density later on led to an increase in the frequency of encounters and gang attacks. However, data suggest that while the frequency of intergroup encounters and gang attack are positively correlated, they appear independent from population density.

Back in 1993-1994, the intergroup encounter rate was 0.18 encounters/day [2], and the population density was 3.9 groups/km$^2$, 68.7 individuals/km$^2$ [3]. The population in the area decreased sharply afterwards ( In 1999: 3.6 groups/km$^2$, 32.4 individuals/km$^2$[4] and then recovered close to the values in 1993-1994 (2009-2010: 44.9 individuals/km$^2$, [5], 2011: 4.3 groups/km$^2$, 61.5 individuals/km$^2$; [4]). The rate of encounters and gang attacks, conversely, seems to have increased steadily over the years (**Table S2**).

| Group | Period | IGE/day | GA/day | No. Observation days |
|---|---|---|---|---|
| **PB/PB1/PB1A** | 2006-2010[1] | 0.46 | 0.002 | 622 |
| | 2015-2016[2] | 1.09 | 0.006 | 170 |
| **R1** | 2006-2010[1] | 1.03 | 0.012 | 1079 |
| | 2015-2016[2] | 1.67 | 0.054 | 129 |
| **R2** | 2006-2010[1] | 0.35 | 0.004 | 804 |
| | 2015-2016[2] | 0.45 | 0 | 31 |

**Table S2. Rate of intergroup encounters (IGE) and intergroup gang attacks (GA) recorded on three groups of crested macaques from Tangkoko National Park in two different periods.** [1]Unpublished data from D.Kerhoas, used in Kerhoas et al. (2014). [2]Unpublished data from L.Martínez-Íñigo, used in [1]

However, we need to be cautious with these conclusions due to the numerous sources of bias. These include a different methodology to estimate population density, differences in the definition of intergroup encounter, and research focus of each period. For example, encounters were usually routinely until 2015-2016, when a research team focused on them. This might explain why the highest intergroup encounter rates were recorded then.

Kinnaird and O'Brien (2000) found a trend towards an increasing number of encounters with an increasing amount of defensible (i.e., patchy) food in the diet of the crested macaques. If the food has become patchier with the years, that might explain the lower population density and more groups with fewer individuals, as well as more frequent intergroup encounters.
